# Potpourri: An Epistasis Test Prioritization Algorithm via Diverse SNP Selection

**DOI:** 10.1101/830216

**Authors:** Gizem Caylak, Oznur Tastan, A. Ercument Cicek

## Abstract

Genome-wide association studies explain a fraction of the underlying heritability of genetic diseases. Investigating epistatic interactions between two or more loci help closing this gap. Unfortunately, sheer number of loci combinations to process and hypotheses to test prohibit the process both computationally and statistically. Epistasis test prioritization algorithms rank likely-epistatic SNP pairs to limit the number of tests. Yet, they still suffer from very low precision. It was shown in the literature that selecting SNPs that are individually correlated with the phenotype and also diverse with respect to genomic location, leads to better phenotype prediction due to genetic complementation. Here, we propose that an algorithm that pairs SNPs from such diverse regions and ranks them can improve prediction power. We propose an epistasis test prioritization algorithm which optimizes a submodular set function to select a *diverse and complementary* set of genomic regions that span the underlying genome. SNP pairs from these regions are then further ranked w.r.t. their co-coverage of the case cohort. We compare our algorithm with the state-of-the-art on three GWAS and show that (i) we substantially improve precision (from 0.003 to 0.652) while maintaining the significance of selected pairs, (ii) decrease the number of tests by 25 folds, and (iii) decrease the runtime by 4 folds. We also show that promoting SNPs from regulatory/coding regions improves the performance (up to 0.8). Potpourri is available at http:/ciceklab.cs.bilkent.edu.tr/potpourri.

## 1 Introduction

Genome-wide association studies (GWAS) have been an important tool for susfaceptibility gene discovery in genetic disorders (1; 2; 3). Analyzing single loci associations have provided many valuable insights but they alone do not account for the full heritability (4). Investigating the interplay among multiple loci has helped to bridge the missing heritability gap. Such interactions between two or more loci is called epistasis and it has a major role in complex genetic traits such as cancer (5) and neurodevelopmental disorders (6).

Exhaustive identification of interacting loci, even just pairs, is potentially intractable for large GWAS (7). Moreover, such an approach lacks statistical power due to multiple hypothesis testing. Several methods have been developed to circumvent these problems. TEAM and BOOST are exhaustive models which exploit data structures and efficient data representation to improve the brute force performance (8; 9). However, these methods still perform many tests and do not scale for large tasks. For instance, BOOST reports a runtime of 60 hours for 360k SNPs. Another approach is to reduce the search space by filtering pairs based on a type of statistical threshold. Examples include SNPHarvester (10), SNPRuler (11) and IBBFS (12). Despite their advantages, these methods mostly do not follow a biological reasoning and tend to detect interactions that are in linkage disequilibrium (LD) as noted in Piriyapongsa *et al.* (13). On another track, incorporating biological background and testing the SNP pairs that are annotated has also proven useful (14; 15; 16; 17; 18). Yet, this approach requires most SNPs to be discarded as many are quite far away from any gene to be associated. Moreover, this introduces a literature bias in the selections of the algorithms.

A rather more popular approach is to prioritize the tests to be performed rather than discarding pairs from the search space and controlling for Type-I error. In this approach, the user can keep performing tests in the order specified by the algorithm until a desired number of significant pairs are found. The idea is to provide the user with a manageable number of true positives (statistically significant epistatic pairs) while minimizing the number of tests to ensure statistical power. The first algorithm of this kind is iLOCi (13), which ranks SNP pairs by performing a dependence test and avoiding pairs that are unrelated to disease but might seem related due to LD. This work was followed by Ayati and Koyuturk who proposed testing pairs of SNPs in *population covering locus sets - POCOs* (19). First, the algorithm greedily selects multiple exclusive groups of SNPs that *cover* all affected individuals. That is, each case sample has to have at least one SNP in each POCO. Epistasis tests then are performed across POCOs with the hope that independent coverage of the cases will lead different POCOs to include complementary and epistatic SNPs (19**?**). Finally, Cowman and Koyuturk, introduced the LINDEN algorithm (7). First, in a bottom-up fashion, the method generates SNP trees on greedily selected genomic regions (LD forest). Each node represents the genotypes of cases and controls for the SNPs in that node. Then, by comparing the roots of these trees, it decides if this pair of regions is a promising candidate for epistasis test. Nodes in lower levels are continued to be checked and leaf pairs (individual SNPs) are tested only if the estimation at higher levels is promising. LINDEN was shown to achieve the state of the art results. Despite using various heuristics, all methods still have high false discovery rates. For instance, the FDR of LINDEN ranges from 0.96 to 0.998 on three GWAS from Wellcome Trust Case Control Consortium (WTCCC) - the ratio of significant pairs to the number of detected (reciprocally) significant epistatic pairs (7).

Linkage disequilibrium is an important source of information for epistasis prioritization algorithms. Two SNPs that appear to be interacting statistically, might not be biologically meaningful if they are on the same haplotype block (20). For this reason, all three methods mentioned above focus on such regions and aim at avoiding testing pairs that are located in close vicinity of each other. In an orthogonal study, Yilmaz et al. propose a feature (SNP) selection algorithm which avoids LD for better phenotype prediction (21). Authors show that while looking for a small set of loci (i.e., 100) that is the most predictive of a continuous phenotype, selecting SNPs that are far away from each other, results in better predictive power. This method, SPADIS, is designed for feature selection for multiple regression. As the SNP set it generates contains diverse and complementary SNPs, it results in better *R*^2^ values by covering more biological functions.

Inspired by this idea, we conjecture that selecting pairs of SNPs from genomic regions that (i) harbor individually informative SNPs, and (ii) are diverse in terms of genomic location would avoid LD better and yield more functionally complementing and more epistatic SNP pairs compared to the current state of the art since no other algorithm exploits this information. We propose a new method that for the first time incorporates the genome location diversity with the population coverage density. Specifically, our proposed method, *Potpourri*, maximizes a submodular set function to select a set of genomic regions (i) that include SNPs which are individually predictive of the cases, and (ii) that are distant from each other on the underlying genome. Epistasis tests are performed for pairs across these regions, such that pairs that densely co-cover the case cohort are given priority.

We validate our hypothesis and show that Potpourri is able to detect statistically significant and biologically meaningful epistatic SNP pairs. We perform extensive tests on three Wellcome Trust Case Control Consortium (WTCCC) GWA studies and compare our method with the state-of-the-art LINDEN algorithm. First, we *guide* LINDEN by pruning its search space using Potpourri-selected-SNPs to show that (i) it is possible to significantly improve the precision (from 0.003 up to 0.302) and (ii) that our diversification approach is sound. Then, we show that the ranking of the diverse SNPs by the co-coverage of the case cohort further improves the prediction power and the precision (up to 0.652 in the selected setting). Potpourri drastically reduces the number of hypothesis tests to perform (from ∼ 380*k* to ∼ 15*k*), and yet is still able to detect more epistatic pairs with similar significance levels in all there GWA studies considered. The running time is also cut by 4 folds in the selected settings. Another problem with the current techniques is the biological interpretation of the obtained epistatic pairs. Once the most significant SNP pairs returned are in the non-coding regions and are too isolated to be associated with any gene, the user can hardly make sense of such a result despite statistical significance. We investigate the advantage of promotion of SNPs falling into regulatory and non-coding regions for testing and propose three techniques. We show that these techniques further improve the precision (up to 0.8) with similar number of epistatic pairs detected. Finally, we investigate the biological meaning of the detected SNP pairs. We find (i) a SNP pair which supports the hypothesis of a shared genetic architecture between T2D and chronic kidney disease; and (ii) a pair which suggests new candidate risk gene (*NPW*) for HT which has loose indications only in rat studies. Potpourri is available at http://ciceklab.cs.bilkent.edu.tr/potpourri.

## 2 Methods

### 2.1 Notation

A GWAS dataset consists of genotypes of a set of samples *S* who are associated with a binary phenotype: *Case* or *Control*. Let *f* (*s*) be an indicator function that corresponds to phenotype of a sample *s* ∈ *S*, i.e.: *f* (*s*) = 1, if *s* is a case sample, and *f* (*s*) = 0, otherwise. Function *h* which represents the genotype of sample *s* ∈ *S* at locus *v* ∈ *V* is encoded as:

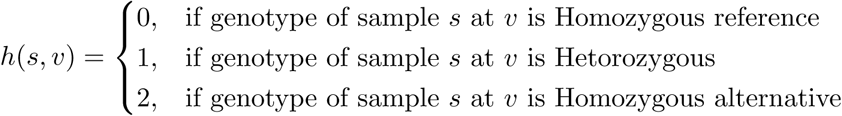

One can generate an undirected SNP-SNP network *G*(*V*′, *E*), where *V*′ ⊆ *V* and *v*_*i*_ ∈ *V*′ if ∃*s* ∈ *S* where *h*(*s, v*_*i*_) ≠ 0. 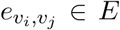 where *v*_*i*_, *v*_*j*_ ∈ *V* and they are *related*. The definition of *relatedness* might change with respect to the application and the biological definition. In our setting, two SNPs are related if they are neighbors on the underlying genome.

### 2.2 Selection of Diverse and Informative SNP Regions

Regions harboring informative SNPs (ones that are correlated with the phenotype) and also far away on the underlying genome with respect to a SNP-SNP network are likely to yield a diverse and explanatory SNP set without introducing a literature bias. Our hypothesis is that those selected SNPs are likely to be epistatic because SNPs selected using this technique leads to better phenotype prediction due to better genomic complementation (21). Hence, picking pairs from such a set should yield highly epistatic SNP pairs and is well situated for epistasis test prioritization. In our application the goal is not to predict the phenotype. Our SNP set selection approach differs from Yilmaz *et al.* in the calculation of the SKAT scores (22) as in epistatis test prioritization we are analyzing a GWA study in which the phenotype is dichotomous (the term 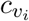 in Eq. 1). Moreover, we introduce an hyper-parameter which can assign extra artificial price to SNPs that fall in regulatory or coding regions to give priority to SNPs which are more likely to have functional effect (the term 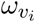 in Eq. 1). Also, node prizes (i.e., SKAT scores) are normalized by the set size to be able to compare performances of selected sets of different sizes (division of 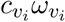 by *k* in Eq. 1).

First, for every SNP *v*_*i*_, a score 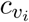 is calculated using SKAT method which works by regressing phenotype on the covariates using a flexible semiparametric linear model (22). Instead of directly associating genotypes of the variants with the phenotype, SKAT uses a nonparametric function of the genotypes that is possibly contained in a vector space generated by a positive semi-definite kernel function. Given a user specified number SNPs to select (*k*), the second step selects a subset of loci *EP* ⊂ *V*′ such that it maximizes the sum of selected SNP scores while penalizing SNPs that are in close vicinity of each other. In particular the function *F* as shown in Equation 1 is maximized:

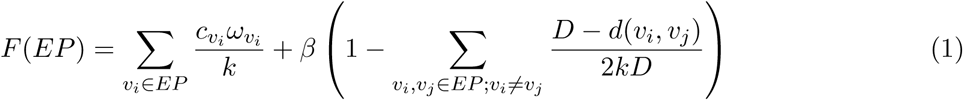

In this equation, *D* ∈ ℝ_*>*0_ is a parameter that sets the upper limit on the distance that two SNPs are considered *close. d*(*v*_*i*_, *v*_*j*_) is the distance between vertices *v*_*i*_, *v*_*j*_ ∈ *EP* on graph *G*. In this application function *d*(*v*_*i*_, *v*_*j*_) denotes the shortest path distance between *v*_*i*_, *v*_*j*_ only if *d*(*v*_*i*_, *v*_*j*_) ≤ *D*, otherwise *d*(*v*_*i*_, *v*_*j*_) = *D* to cancel out the penalty term. The parameter *β* ∈ ℝ_≥0_ adjusts the relative magnitudes of the prices and the penalties. Finally, the parameter 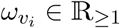 rewards *v*_*i*_ ∈ *V*′ if it falls into a regulatory or coding region of interest. If this information is not taken into consideration 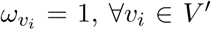 - see Subsection 2.3.3. The subset *EP*\* that maximizes *F* is selected as the seed set for epistasis prioritization as shown in Equation 2.

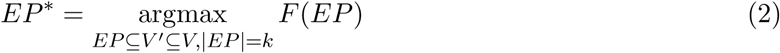

Given the set of SNPs *V*′, the above-mentioned function *F*: 2^*V′*^ → ℝ is a submodular function (see Supplementary Text 1.1 for the proof). Submodular optimization is NP-hard. However, the greedy algorithm given in Supplementary Text 1.2, which basically iteratively adds the next *best* SNP to the set that maximizes *F* at each step, ensures a 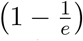 factor approximation to the optimum solution (23). This algorithm requires the submodular set function to be also monotone non-decreasing and non-negative, which are proven in Supplementary Text 1.3.

Note that using only the SNPs in *EP*\* would restrict the epistasis testing to SNPs which are individually correlated with the trait. We rather use those as anchors to detect genomic regions of interest. Let *R* be a set of consecutive SNPs that fall into a region in the genome. After *EP*\* is set, for every SNP *v*_*i*_ ∈ *EP*\*, a region *R*_*i*_ is formed such that it includes, *v*_*i*_ and *m* upstream and *m* downstream SNPs of *v*_*i*_. Thus, |*R*_*i*_| = 2*m* + 1. **R** is the set of sets (regions) *R*_*i*_, ∀*v*_*i*_ ∈ *EP*\*. Note that 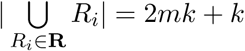.

### 2.3 Prioritization of the Tests

The region set **R** generates a pruned search space for finding likely epistatic SNP pairs. Within region tests are avoided in order to avoid LD. That is, SNPs *v*_*i*_ and *v*_*j*_ are not tested if both *v*_*i*_ ∈ *R*_*x*_ and *v*_*j*_ ∈ *R*_*x*_ such that *R*_*x*_ ∈ **R**. Still, the number of possible tests is 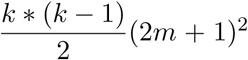 and a prioritization scheme is needed to rank candidate pairs for testing. We employ two strategies. The first method uses our SNP selection strategy to prune the search space for LINDEN to show that the selected regions are good candidates for epistasis testing - see Section 2.3.1. Instead of guiding LINDEN, in the second approach, we introduce a new method that aims at testing the pairs which co-occur in the case cohort - see Section 2.3.2.

#### 2.3.1 Guiding LINDEN for Ranking SNP Pairs

In this ranking strategy, we use LINDEN to rank the pairs selected in **R**. That is, we use Potpourri selected SNPs to guide LINDEN. Thus, in this strategy, Potpourri acts as a preprocessing step for LINDEN to prune its large search space. Doing so, we test if our diverse SNP selection scheme is sound and whether it improves the performance of LINDEN.

LINDEN is input with only SNPs that fall into the regions in **R**. Then, the algorithm is run as described in Cowman and Koyuturk. That is, the algorithm forms LD trees over greedily selected regions. The leaves of these trees represent actual SNPS each of which are represented by a sample genotype vector (i.e., the genotype of this SNP for every sample). The tree is constructed in bottom-up fashion and nodes are merged to genertate higher levels. During this process, sample genotype vectors are merged and ambiguous indices (i.e., samples with different genotypes) are assigned a *NIL* value. This merging step continues until a threshold *d* is met that denotes the fraction of *NIL* values allowed. This is a dynamically adjusted threshold goes up with the number of iterations. Then, the tree pairs are tested with respect to the sample genotype vectors starting from the root, ignoring the *NIL* values. Lower levels are tested only if the significance of the chi-squared test meets a certain threshold. In parallel with our hypothesis on diversification of the regions, we prohibit LINDEN to merge SNPs from different regions. Thus, the tests are performed across regions in **R**.

#### 2.3.2 Population Co-covering For Ranking SNP Pairs

We propose a new strategy which aims at maximizing the population coverage of co-occuring SNPs. Population cover for epistasis test prioritization was first proposed in Ayati *et al.* which selects multiple exclusive groups of SNPs (*POCO*s) that *covers* the case cohort (19**?**). That is, the union of the samples with the SNPs in a POCO should be equal to the case cohort. Epistasis tests are performed across POCOs and the idea is that the independent coverage of the cases across POCOs will result in detecting complementary SNPs. We take a different approach in terms of covering the population. We would like the SNP pairs to be tested to co-cover the population. This is intuitive; the diverse selection step described in Section 2.2 finds complementing regions and avoids LD, but for a SNP pair to be epistatic they also need to be observed together in cases. More formally, let *p*(*v*_*x*_, *v*_*y*_) be a function that scores the SNP pair *v*_*x*_, *v*_*y*_ ∈ *V*′ for testing. Given three SNPs *v*_*x*_, *v*_*y*_ and *v*_*z*_ ∈ *V*′, *p*(*v*_*x*_, *v*_*y*_) *> p*(*v*_*x*_, *v*_*z*_) if and only if *v*_*x*_ is observed more frequently with *v*_*y*_ in cases as compared to *v*_*z*_. *p* is formally defined as follows:

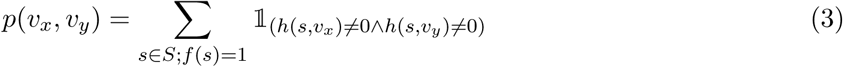

Potpourri computes the pairwise population co-covering of all SNP pairs (*SNP*_*i*_, *SNP*_*j*_) such that *SNP*_*i*_ ∈ *R*_*x*_, *SNP*_*j*_ ∈ *R*_*y*_, ∀*R*_*x*_, *R*_*y*_ ∈ **R**, *x* ≠ *y* and *i* ≠ *j*. Testing is then performed in the descending population co-cover order. The algorithm is restricted to test top *v* pairs among all region pairs in **R**. See Section 2.3.3 to for details of prioritization when regulatory/coding regions are considered.

#### 2.3.3 Promoting SNPs in Regulatory and Coding Regions

SNPs can alter gene expression and the downstream function depending on their genomic location. Those that fall on to regulatory regions can affect mRNA levels and those that fall onto coding regions can alter the structure and function of the the protein. Since such SNPs are likely to alter the function and more likely to be related to the phenotype, we conjecture that we can find more statistically significant and biologically meaningful SNP pairs via promoting mutations in regulatory and coding regions. While this might introduce a literature bias, for a life scientist who would like to narrow down the search space using functional regions this might be plausible and investigate the usefulness of this approach. However, one should not totally disregard the unannotated parts of the genome as most of the variation exists in such regions. Thus, we seek a balance.

We employ 3 techniques to *promote* regulatory/coding variants. In the first approach (Potpourri RC1), we assign an artificial prize to SNPs in the diverse SNP selection phase of the algorithm as described in Section 2.2. This approach favors the regions in *R* to be regulatory/coding regions. Then, the pairs are tested with respect to the population co-cover ranking. The second approach (Potpourri RC2), uses 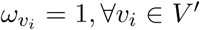. That is, the SNP selection step does not promote selection of coding/regulatory SNPs. However, among the selected SNPs, the prioritization step prefers pairs such that at least one SNP falls into regulatory/coding regions. Then, these pairs are ranked with respect to the population co-cover as described in Section 2.3.2. Note that after all such pairs are tested, the algorithm test remaining pairs with respect to their population co-cover ranking. The final strategy (Potpourri RC3) combines both RC1 (promoting selection of SNPs in regulatory/coding regions) and RC2 (first testing pairs with at least one SNP in regulatory/coding regions) strategies.

## 3 Results

### 3.1 Datasets

In order to benchmark Potpourri, we used three GWAS datasets obtained from Wellcome Trust Case Control Consortium (WTCCC): (i) Type 2 diabetes (T2D), (ii) Hypertension (HT), and (iii) Bipolar disorder (BD). We use the 1958 Birth (58C) cohort as control for all datasets (24). Please see the details of preprocessing and quality control steps taken in Supplementary Text 1.4.

We used the following resources to obtain the regulatory and coding region information. We obtained the distant-acting transcriptional enhancer dataset from VISTA Enhancer Browser (25). VISTA enhancer dataset contains 1912 human noncoding fragments with gene enhancer activity. Transcription start sites (TSSs) and coding region coordinates were obtained from UCSC Genome Browser (26). We chose Ensembl Genes as gene annotation track (27) and the March 2006 NCBI36/hg18 assembly of the human genome which matches the WTCCC datasets. The number and types of genes obtained from the Ensembl dataset are given in Supplementary Table 2. We defined 1 Kb downstream and upstream of the TSSs as a regulatory region. The coding region for a gene begins from the first base of the first exon and continues to the last base of the last exon.

Potpourri operates on a SNP-SNP interaction network. In this study, we used the genomic sequence (GS) network as defined in Azencott *et al.* (28). In this network, SNPs are connected if they are adjacent on the genome. This was the network of choice as it shown to provide the best regression performance in Yilmaz *et al.*.

### 3.2 Experimental Setup

We follow the experimental procedure in Yilmaz et al. and Azencott et al. for the SNP selection step (21; 28). The parameters were selected using a nested 10-fold cross validation. First, the distance parameter *D* was selected via a line search in among 7 values (log-scale) between 1 and *D*_*max*_ (a value for which the distance penalty for all SNPs in the selected set is 0). *D* value that maximizes the L2-regularized logistic regression performance was picked. Then, 16 *β* values between 10^−4^ and a maximum *β* = 2*kD*_*max*_ were tried. Again, the *β* value that maximizes the classification performance was picked. We experimented with the following *k* values: 500, 750, 1000, 1500, and 2000. Overall, best results were obtained when *k* = 750 and it was set as the default parameter setting. Then, we added *m* = 9 upstream and *m* = 9 downstream neighbors SNPs of the for further coverage analysis. Unless otherwise stated, we set the *w*_*i*_ = 1, ∀*v*_*i*_ ∈ *V*′. Once the regulatory and coding regions were considered by the diverse SNP selection part, *w*_*i*_ *>* 1, ∀*i* ∈ *RC* where *RC* contains SNPs that fall into regulatory and coding regions. We experimented with the three *ω* values: 1 + 10^−0.5^, 1 + 10^−0.25^, and 2.

Unless otherwise stated, we used the suggested settings for LINDEN as described in Cowman and Koyuturk: *d* = .45 and *b* = 10. When guiding LINDEN with our approach, we used default parameters for LINDEN. The only exception was that, we limited the extent of LD in the first two iterations of the merging procedure of LINDEN as explained in Section 2.3.1. That is, LINDEN’s merging step could merge SNPs form trees only within selected regions that contain of 2*m* + 1 SNPs (*m* = 9 in our experiments). We used the above-mentioned parameter selection techniques for Potpourri. For the population co-cover, we performed epistasis test for the top *v* = 10 SNP pairs among regions in **R**.

To quantify the performance of the proposed algorithms, we used precision as the evaluation metric in which true positives (TP) refer to the reciprocally significant epistatic pairs that pass the Bonferroni-adjusted statistical significance threshold. False positives (FP) refer to failed tests: the reciprocally significant epistatic pairs that fail to pass the aforementioned threshold. Note that, in the epistasis test prioritization context, a false positive does not mean a false rejection of the null hypothesis and falsely concluding that the pair has an epistatic interaction. Rather, it refers to a prioritized pair for testing that could not pass the corrected statistical significance threshold.

We use the definition of *reciprocally significant epistatic pair* from Cowman and Koyuturk. As the authors argue, most epistatic pairs are dominated by some hub SNPs and this leads to detection of redundant pairs. On the other hand, SNPs in a reciprocal pair are the most epistatic partner for each other. We also use this definition to measure the performance of our algorithm. We set the significance level as 10% throughout experiments and adjust the significance level using the Bonferroni correction based on the number of test performed by each approach. Epistasis testing is performed via a chi-squared test on a 9×2 contingency table of all genotype combinations between cases and controls for a selected SNP pair (df = 8) as also done by Cowman and Koyuturk. All tests are performed on an Intel(R) Xeon(R) E5-2650 v3 Ten-Core Haswell Processor (2.3GHz 25M 9.6GT/s QPI). 251 GB RAM is used in the parallel setting.

### 3.3 Guiding LINDEN with Diverse and Informative Regions Improves Precision

We quantified the improvement in precision when LINDEN is guided by Potpourri-selected regions. First, we ran LINDEN on all datasets and it detected 1786, 885, and 1022 reciprocally significant epistatic pairs for T2D, BD, and HT datasets, respectively. Only 5, 35, and 2 pairs were statistically significant at the 10% level after Bonferroni correction, respectively. These numbers correspond to precision values of 0.003, 0.04, and 0.002, respectively. The selected SNP Pairs by LINDEN are listed in Supplementary Tables 27 - 29, for T2D, BD and HT datasets, respectively. These results set our baseline. Then, we ran Potpourri on these datasets with top *k* SNPs, where *k* = 500, 750, 1000, 1500 and 2000. We obtained 5 **R** sets. We then *guide* LINDEN with these regions and ran it as explained in Section 2.3.1.

Complete results are shown in Table 1, Supplementary Table 3, and Supplementary Table 4 for T2D, BD and HT datasets, respectively. The guidance of Potpourri improves the precision substantially, from 0.003 to 0.3 when *k* = 750 in the T2D dataset (up to .421 when *k* = 500). This is achieved by drastically reducing the number of false positives, and also increasing the number of true positives. Our pipeline outperforms LINDEN for all *k* values on all datasets, but we observe that the ideal *k* values are 500, 750 and 1000. Similar precision increases are also observed for BD and HT datasets. Statistics on the number of tests performed and trees formed by LINDEN and Potpourri-guided LINDEN are shown in Supplementary Table 6.

**Table 1:** Results for T2D dataset that compares LINDEN, Potpourri-guided LINDEN and Potpourri (with population co-cover). Number of pairs reported is the total number of reciprocally significant pairs returned by each method for varying number of selected SNPs. The number in parentheses denotes the significant pairs passing significance threshold (0.1) after Bonferroni correction based on the number of tests performed by each method. Bold denotes the best result for a given *k* value. The table shows that the guidance of Potpourri substantially improves the precision of LINDEN. For all considered *k* values, Potpourri provides the best precision values.

| Method |  | # Tested Loci | # Reported (Rec. Sig.) | Precision |
| --- | --- | --- | --- | --- |
| **LINDEN** |  | 378016 | 1786 (5) | 0.003 |
| **Potpourri guided LINDEN** | $k = 500$ | 9250 | 19 (8) | 0.421 |
| | $k = 750$ | 14146 | 43 (13) | 0.302 |
| | $k = 1000$ | 18943 | 55 (21) | 0.382 |
| | $k = 1500$ | 28014 | 79 (18) | 0.228 |
| | $k = 2000$ | 37927 | 95 (11) | 0.116 |
| **Potpourri** | $k = 500$ | 9250 | **12 (8)** | **0.667** |
| | $k = 750$ | 14145 | **23 (15)** | **0.652** |
| | $k = 1000$ | 18943 | **31 (20)** | **0.645** |
| | $k = 1500$ | 28014 | **50 (21)** | **0.420** |
| | $k = 2000$ | 37927 | **74 (18)** | **0.243** |

The significance levels of each performed epsistasis test by LINDEN and Potpourri-guided LINDEN are shown in Figure 1, Supplementary Figure 1, and Supplementary Figure 2 for the T2D, BD and HT datasets, respectively. The green lines denote the significance level (0.1) to be passed for each approach (*k* = 750) after Bonferroni correction. Each point represents a SNP pair and the ones below threshold are false positives and the ones above are true positives (reciprocally significant epistatic SNP pairs). It is clear that the pipeline drastically reduces the number of false positives while increasing the number true positives. Also, we can observe the importance of the number of tests performed by looking at the difference between Bonferroni corrected significance thresholds. Since Potpourri provides a pruned search space for LINDEN by eliminating SNPs that are most likely irrelevant to the trait, it also reduces number of tests that will be performed during epistasis test. Indirectly, it eliminates the negative effect of multiple hypothesis testing which reduces the statistical power. As seen in the figures, due to low Bonferroni threshold, the pipeline is able to discover more TPs compared to LINDEN. We also show that our pipeline not only minimizes the number of false positives but is also able to maintain the significance level of the returned pairs. While LINDEN’s top SNP pair is more significant, *guided* LINDEN’s top SNP pairs still stand out in terms of their p-values. We compared the sets of SNP pairs detected by LINDEN and Potpourri-guided LINDEN and found that there the former is not a superset of the latter. Potpourri enforces LINDEN to form trees on its selected regions of 2*m* + 1 SNPs and tests are performed across these trees. LINDEN follows a greedy approach to form trees in a bottom-up fashion. Thus, the trees it forms are potentially much different (e.g., deeper, covering more loci) than the guided version and the returned pairs are different.

**Figure 1:**
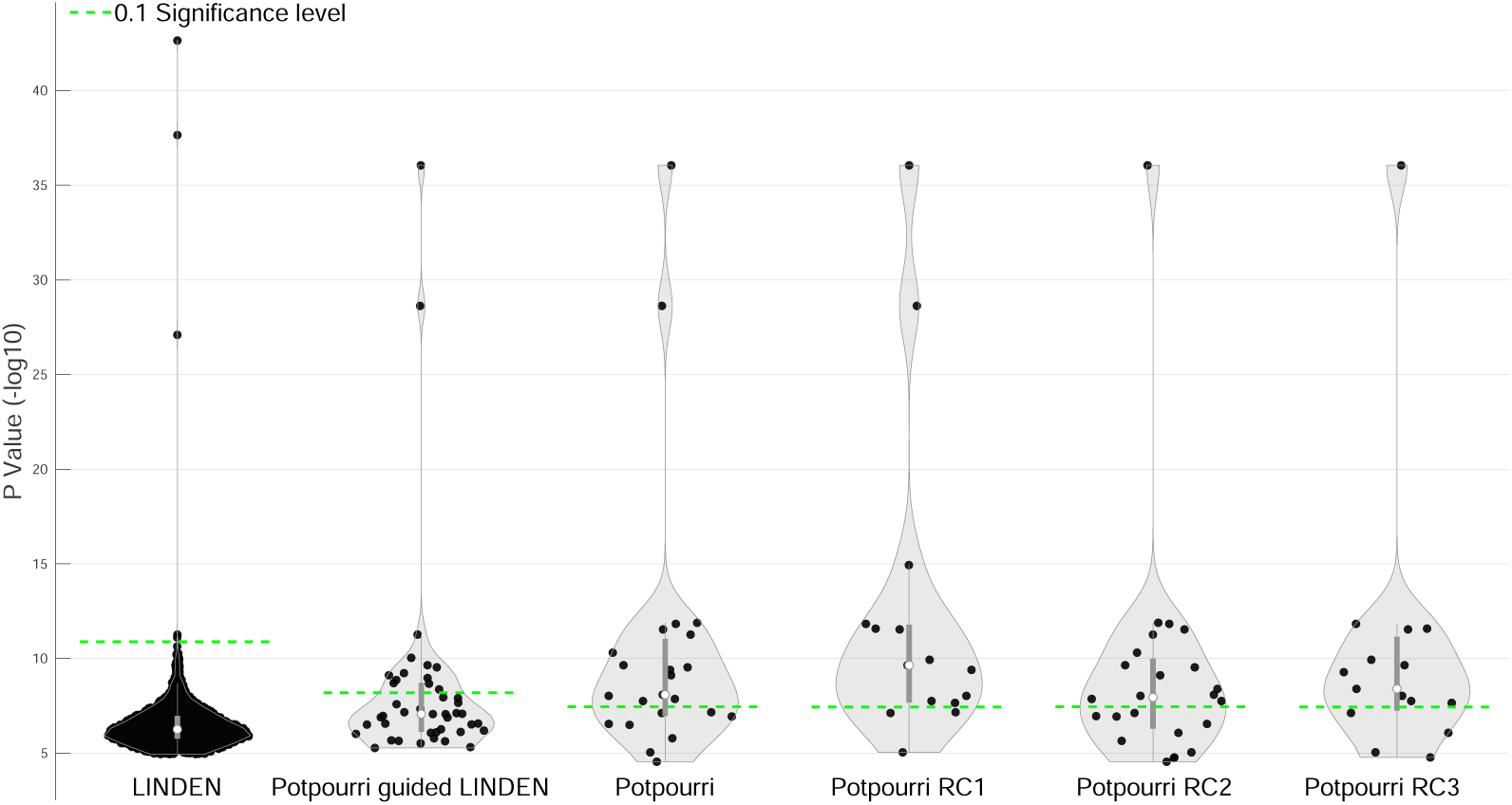
On the T2D dataset, this figure compares the p-values of the selected SNP pairs by the following methods: (i) LINDEN, (ii) Potpourri *guided* LINDEN, (iii) Potpourri; and then also the variants of Potpourri which further promotes SNP pairs in regulatory and coding regions: (iv) Potpourri RC1, (v) Potpourri RC2, and finally (vi) Potpourri RC3. We show the significance levels (y-axis) of each reported pair (dots) given the Bonferroni-corrected significance threshold (0.1, green dashed lines). X – axis is just randomly assigned values to pairs for better visualization. Potpourri is run with *k* = 750 for all 5 related subplots. For RC1 and RC3 strategies *ω* is set to 1.31623, for Potpourri and RC2, it is set to 1. LINDEN is run with default parameters. For Potpourri *guided* LINDEN, tree formation is restricted to be done on distinct regions in Potpourri-selected regions **R** as described in Section 2.3.1.

In short, by just providing a guidance to LINDEN using Potpourri’s diverse and informative SNP region selection, we were able to improve the performance of the state-of-the-art with substantially smaller number of tests performed. Next, we provide the results of the complete Potpourri pipeline which uses population co-cover strategy for prioritization, instead of LINDEN’s tree based strategy.

### 3.4 Comparison of Potpourri with the State-of-the-art

In this section, we evaluate the performance of *the* Potpourri pipeline with population co-covering technique. Again, we compare the performance with the LINDEN algorithm. Table 1 shows the results of the algorithm for different *k* values on the T2D dataset. We observe that population co-cover ranking strategy results in even further performance improvements. The precision is moved up to 0.652, as 15 out of 23 reciprocally significant pairs passes the Bonferonni-corrected threshold (*k* = 750). See Supplementary Tables 3 and 4 for the results on BD and HT datasets, respectively, which follow the same pattern.

LINDEN performs 7,629,272,394 tests (378,016 tests for leaves) as opposed to 14,146 tests performed by Potpourri. This sets a much more conservative significance threshold. One could argue that the precision gain is only due to the reduced significance threshold. However, Figure 1 shows that it is not the case. Third panel shows the significance levels of Potpourri-selected SNP pairs on the T2D dataset. We see that 6 out of 15 Potpourri-selected reciprocally significant pairs are significant even when the Bonferonni-corrected threshold of LINDEN is considered (dashed green line on the left-most panel which is far more conservative). Note that LINDEN only detects 5 reciprocally significant pairs. We also observe that the number of false positives are even further decreased compared to Potpourri-guided LINDEN. See Supplementary Figures 1 and 2 for the results on BD and HT datasets, respectively, which again show a similar pattern.

We also checked if adjusting LINDEN’s parameters to make it more conservative would improve the precision. Increasing *d* to .9 and to .99 enforced it to perform a smaller number of tests but still too many to get close to Potpourri’s efficiency, as shown in Supplementary Table 5. We checked if we would get better results by testing only SPADIS selected SNPs. For each dataset, we performed pairwise epistasis tests on all SPADIS-selected SNPs (*k* = 750). In each dataset, only 1 reciprocally significant pair is detected. Finally, we show that in the suggested setting (*k* = 750) the time requirement is decreased by 4 folds for all datasets as shown in Supplementary Tables 30 - 32.

### 3.5 Promoting Regulatory and Coding Regions Improves Precision

In this section, we investigate the advantage of promoting coding and regulatory regions in the Potpourri pipeline. For the T2D dataset, Table 2 compares the performance of original Potpourri pipeline with the three variants (RC1, RC2 and RC3) that promote selection of SNPs from coding and regulatory regions as described in Section 2.3.3. Results show that in the suggested setting (*k* = 750) RC1 technique can increase the precision up to 0.8 and the original Potpourri pipeline cannot achieve a better result in any of the *k* values (see Table 2). Last 3 panels in Figure 1 show the significance levels of tested pairs. For the RC1 technique, we see that while we keep the most significant pairs detected by the original pipeline, we also discover new significant pairs (i.e., *p <* 10^−15^). Moreover, the 2 out of 3 pairs that failed to pass the significance threshold are borderline. 7 out of 15 reciprocally significant pairs are also significant with respect to LINDEN’s stringent significance threshold while LINDEN was able to detect only 5 such pairs and Potpourri was able to detect 6. Supplementary Tables 11 and 14 provide similar results for the BD and HT datasets, respectively. The margin of improvement is relatively low in the HT dataset (up to ∼0.2). Supplementary Figures 1 and 2 compare the significance levels of the detected pairs for the BD and HT datasets, respectively.

**Table 2:** Comparison of the Potpourri results with three strategies (RC1, RC2,and RC3) to promote regulatory and coding regions on the T2D dataset. For RC1 and RC3 *ω* is set to 1.31623. Number of pairs reported is the total number of reciprocally significant pairs. The number in parentheses denotes the significant pairs passing significance threshold (0.1) after Bonferroni correction based on the number of tests performed by each method. Results show that promoting regulatory and coding regions improves Potpourri results substantially in all strategies, RC1 performing the best. Bold denotes the best result for a given *k* value.

| Method |  | # Tested Loci | # Reported (Rec. Sig.) | Precision |
| --- | --- | --- | --- | --- |
| Potpourri | $k = 500$ | 9250 | 12 (8) | 0.667 |
| | $k = 750$ | 14145 | 23 (15) | 0.652 |
| | $k = 1000$ | 18943 | 31 (20) | 0.645 |
| Potpourri RC1 | $k = 500$ | 9489 | **17 (13)** | **0.765** |
| | $k = 750$ | 13727 | **15 (12)** | **0.800** |
| | $k = 1000$ | 15500 | 21 (12) | 0.571 |
| Potpourri RC2 | $k = 500$ | 9250 | 13 (7) | 0.538 |
| | $k = 750$ | 14145 | 24 (14) | 0.583 |
| | $k = 1000$ | 18943 | **33 (23)** | **0.697** |
| Potpourri RC3 | $k = 500$ | 9489 | 18 (11) | 0.611 |
| | $k = 750$ | 13727 | 15 (11) | 0.733 |
| | $k = 1000$ | 15500 | 20 (13) | 0.650 |

We experimented with three *ω* values and picked the 1.31623 as the best performing and suggested parameter value. It is used to produce above-mentioned figures and tables. See Supplementary Tables 7 - 10, 12, and 13 for all results obtained using other parameter choices. Finally, selected SNP pairs by all Potpourri variant techniques with suggested settings (*k* = 750 and *ω* = 1 or 1.31623) are listed in Supplementary Tables 15 - 26. We also compared the returned reciprocally significant epistatic SNP pairs of LINDEN and Potpourri RC1 (suggested setting) and found no overlaps. The search space is vast and two algorithms have different approaches which results in different result sets.

### 3.6 Novel Epistatic Pairs

One interesting pair discovered by Potpourri is the *rs*12548378 - *rs*10787472 pair that is detected for T2D. The former falls into *TCF7L2* gene and the latter falls into long intergenic non-coding RNA LINC01111. On one hand, *TCF7L2* is a well known type 2 diabetes susceptibility gene, which affects the blood glucose balance by regulating of proglucagon gene expression over the WNT signaling pathway (29). Its other epistatic interactions were also reported in the literature (30). On the other hand, LINC0111 was shown to be related to urine creatinine level in UK Biobank samples (genome-wide significant, *p* = 5.28*E* − 09) (31). Elevelated levels of urine albumin-to-creatinine ratio (UACR) is a biomarker for diabetic nephropathy (DN.) DN occurs as a result of the damage to renal nerves due to high glucose levels in blood (32). While, T2D causes nephropathy over nerve damage, it has also been long debated that two might have overlapping genetic architectures. At least four T2D loci are associated with kidney function in American Indians, and two are related to kidney disease partially independently of T2D (33). Hussain *et al.* reports that a variant in *TCF7L2* causes renal dysfunction but this affect is not independent of T2D in south Indian population (34). Buraczynska *et al.* report that a variant in *TCF7L2* is significantly more frequent in patients with diabetic nephropathy compared to non-diabetic nephropathy (35). Finally, an interaction between *EGFR* and *RXRG* genes was reported which increases the risk of DN in a T2D cohort (36). Thus, this new epistatic interaction further supports the hypothesis of a shared genetic architecture between these two diseases and that T2D related loci can also genetically increase risk for DN.

In the HT data, we detect an epsitatic interaction between *rs*34585560 in *MECOM* gene and *rs*8051877 in NPW gene. MECOM is a transcriptional regulator and is a well-known risk gene for high blood pressure as detected in a large scale GWAS (200k samples) (37). However, not much is known about the mechanism over which *MECOM* affects blood pressure (38). NPW is a G-coupled protein activator which regulates neuroendocrine signals. While its ties to hypertension in humans are loose, Pate *et al.* and Yu *et al.* demonstrate in rats that exogenously applied *NPW* increases mean arterial pressure (39; 40). Thus, this new epistatic pair finding can suggest *NPW* as a new risk gene candidate for HT coupled with the affect of *MECOM*.

## 4 Conclusion

Detecting epistatic interactions is a promising direction for understanding the genetic underpinnings of complex traits, but the sheer number of possible hypotheses to test prohibit brute force techniques. We proposed a new test prioritization technique and showed that selecting individually informative and topologically diverse SNPs in terms of genomic location leads to detecting statistically significant epistatic interactions. Our approach performs favorably compared to the state of the art.

## References

[1] Samani, N. J. et al. Genomewide association analysis of coronary artery disease. New England Journal of Medicine 357, 443–453 (2007).

[2] Easton, D. F. et al. Genome-wide association study identifies novel breast cancer susceptibility loci. Nature 447, 1087 (2007).

[3] Rivas, M. A. et al. Deep resequencing of gwas loci identifies independent rare variants associated with inflammatory bowel disease. Nature genetics 43, 1066 (2011).

[4] Manolio, T. A. et al. Finding the missing heritability of complex diseases. Nature 461, 747 (2009).

[5] Wang, X., Fu, A. Q., McNerney, M. E. & White, K. P. Widespread genetic epistasis among cancer genes. Nature communications 5, 4828 (2014).

[6] Moore, J. H. & Mitchell, K. J. The role of genetic interactions in neurodevelopmental disorders. In The Genetics of Neurodevelopmental Disorders, 69–80 (John Wiley & Sons, Inc Hoboken, NJ, USA, 2015).

[7] Cowman, T. & Koyutürk, M. Prioritizing tests of epistasis through hierarchical representation of genomic redundancies. Nucleic acids research 45, e131–e131 (2017).

[8] Zhang, X., Huang, S., Zou, F. & Wang, W. Team: efficient two-locus epistasis tests in human genome-wide association study. Bioinformatics 26, i217–i227 (2010).

[9] Wan, X. et al. Boost: A fast approach to detecting gene-gene interactions in genome-wide case-control studies. The American Journal of Human Genetics 87, 325–340 (2010).

[10] Yang, C. et al. Snpharvester: a filtering-based approach for detecting epistatic interactions in genome-wide association studies. Bioinformatics 25, 504–511 (2008).

[11] Wan, X. et al. Predictive rule inference for epistatic interaction detection in genome-wide association studies. Bioinformatics 26, 30–37 (2009).

[12] Chuang, L.-Y., Chang, H.-W., Lin, M.-C. & Yang, C.-H. Improved branch and bound algorithm for detecting snp-snp interactions in breast cancer. Journal of clinical bioinformatics 3, 4 (2013).

[13] Piriyapongsa, J. et al. iloci: a snp interaction prioritization technique for detecting epistasis in genome-wide association studies. In BMC genomics, vol. 13, S2 (BioMed Central, 2012).

[14] Mckinney, B. & Pajewski, N. Six degrees of epistasis: statistical network models for gwas. Frontiers in genetics 2, 109 (2012).

[15] Holmans, P. et al. Gene ontology analysis of gwa study data sets provides insights into the biology of bipolar disorder. The American Journal of Human Genetics 85, 13–24 (2009).

[16] Weng, L. et al. Snp-based pathway enrichment analysis for genome-wide association studies. BMC bioinformatics 12, 99 (2011).

[17] Liu, Y. et al. Gene, pathway and network frameworks to identify epistatic interactions of single nucleotide polymorphisms derived from gwas data. BMC systems biology 6, S15 (2012).

[18] Baranzini, S. E. et al. Pathway and network-based analysis of genome-wide association studies in multiple sclerosis. Human molecular genetics 18, 2078–2090 (2009).

[19] Ayati, M. & Koyutürk, M. Prioritization of genomic locus pairs for testing epistasis. In Proceedings of the 5th ACM Conference on Bioinformatics, Computational Biology, and Health Informatics, 240–248 (ACM, 2014).

[20] Cordell, H. J. & Clayton, D. G. Genetic association studies. The Lancet 366, 1121–1131 (2005).

[21] Yilmaz, S., Tastan, O. & Cicek, E. Spadis: An algorithm for selecting predictive and diverse snps in gwas. IEEE/ACM transactions on computational biology and bioinformatics (2019).

[22] Wu, M. C. et al. Rare-variant association testing for sequencing data with the sequence kernel association test. The American Journal of Human Genetics 89, 82–93 (2011).

[23] Nemhauser, G. L., Wolsey, L. A. & Fisher, M. L. An analysis of approximations for maximizing submodular set functions—i. Mathematical Programming 14, 265–294 (1978). URL https://doi.org/10.1007/BF01588971.

[24] Craddock, N. J. et al. Genome-wide association study of cnvs in 16,000 cases of eight common diseases and 3,000 shared controls (2010).

[25] Visel, A., Minovitsky, S., Dubchak, I. & A Pennacchio, L. Vista enhancer browser—a database of tissue-specific human enhancers. nucleic acids res 35:d88-92. Nucleic acids research 35, D88–92 (2007).

[26] Lindblad-Toh, K. et al. A high-resolution map of human evolutionary constraint using 29 mammals. Nature 478, 476 EP – (2011). URL https://doi.org/10.1038/nature10530. Article.

[27] Zerbino, D. R. et al. Ensembl 2018. Nucleic acids research 46, D754–D761 (2017).

[28] Azencott, C.-A. et al. Efficient network-guided multi-locus association mapping with graph cuts. Bioinformatics 29, i171–i179 (2013).

[29] Grant, S. F. et al. Variant of transcription factor 7-like 2 (tcf7l2) gene confers risk of type 2 diabetes. Nature genetics 38, 320 (2006).

[30] An, P. et al. Epistatic interactions of cdkn2b-tcf7l2 for risk of type 2 diabetes and of cdkn2b-jazf1 for triglyceride/high-density lipoprotein ratio longitudinal change: evidence from the framingham heart study. In BMC proceedings, vol. 3, S71 (BioMed Central, 2009).

[31] Zanetti, D. et al. Genetic analyses in uk biobank identifies 78 novel loci associated with urinary biomarkers providing new insights into the biology of kidney function and chronic disease. bioRxiv 315259 (2018).

[32] Bouhairie, V. E. & McGill, J. B. Diabetic kidney disease. Missouri medicine 113, 390 (2016).

[33] Franceschini, N. et al. The association of genetic variants of type 2 diabetes with kidney function. Kidney international 82, 220–225 (2012).

[34] Hussain, H. et al. Tcf7l2 rs7903146 polymorphism and diabetic nephropathy association is not independent of type 2 diabetes—a study in a south indian population and meta-analysis. Endokrynologia Polska 65, 298–305 (2014).

[35] Buraczynska, M. et al. Transcription factor 7-like 2 (tcf7l2) gene polymorphism and clinical phenotype in end-stage renal disease patients. Molecular biology reports 41, 4063–4068 (2014).

[36] Hsieh, C.-H. et al. Analysis of epistasis for diabetic nephropathy among type 2 diabetic patients. Human molecular genetics 15, 2701–2708 (2006).

[37] Ehret, G. B. et al. Genetic variants in novel pathways influence blood pressure and cardiovascular disease risk. Nature 478, 103 (2011).

[38] Coffman, T. M. Under pressure: the search for the essential mechanisms of hypertension. Nature medicine 17, 1402 (2011).

[39] Pate, A. T., Yosten, G. L. & Samson, W. K. Neuropeptide w increases mean arterial pressure as a result of behavioral arousal. American Journal of Physiology-Regulatory, Integrative and Comparative Physiology 305, R804–R810 (2013).

[40] Yu, N. et al. Cardiovascular actions of central neuropeptide w in conscious rats. Regulatory peptides 138, 82–86 (2007).

