## Supplementary Material for "Potpourri: An Epistasis Test Prioritization Algorithm via Diverse SNP Selection"

### Supplementary Material for
### Potpourri: An Epistasis Test Prioritization Algorithm via Diverse SNP Selection

Gizem Caylak, Oznur Tastan, A. Ercument Cicek

#### 1 Supplementary Text

##### 1.1 $F$ is a submodular set function

A submodular function is a set function with diminishing returns property. That is, the increase in the function value decreases as a new item is added. $F$ is a submodular set function as shown next. The marginal gain function $G$ of $F$ when one SNP $v_x \in V'$ is added to the SNP set $EP$ is defined as follows: $G(EP, v_x) = F(EP \cup \{v_x\}) - F(EP)$ where $v_x \notin EP$. Then,

$$\begin{aligned} G(EP, v_x) &= \sum_{v_i \in EP \cup \{v_x\}} \frac{c_i \omega_i}{k} + \beta \left(1 - \sum_{v_i, v_j \in EP \cup \{v_x\}; v_i \neq v_j} \frac{D - d(v_i, v_j)}{2kD}\right) \\ &\quad - \sum_{v_i \in EP} \frac{c_i \omega_i}{k} + \beta \left(1 - \sum_{v_i, v_j \in EP; v_i \neq v_j} \frac{D - d(v_i, v_j)}{2kD}\right) \\ &= \frac{c_{v_x} \omega_{v_x}}{k} + \beta - \frac{\beta}{2kD} \sum_{v_i \in EP} (2D - 2d(v_i, v_x)) \\ &= \frac{c_{v_x} \omega_{v_x}}{k} + \beta - \frac{\beta}{k} \sum_{v_i \in EP} \left(1 - \frac{d(v_i, v_x)}{D}\right) \end{aligned} \tag{1}$$

$F$ is *submodular* if and only if $G(EP, v_x) \geq G(EP', v_x)$ for all sets $EP, EP'$ where $EP \subset EP' \subset V'$, $v_x \in V'$, and $v_x \notin EP'$. Thus, $F$ is submodular if and only if $G(EP, v_x) - G(EP', v_x) \geq 0$.

$$\begin{aligned} G(EP, v_x) - G(EP', v_x) &= \frac{c_{v_x} \omega_{v_x}}{k} + \beta - \frac{\beta}{k} \sum_{v_i \in EP} \left(1 - \frac{d(v_i, v_x)}{D}\right) \\ &\quad - \frac{c_{v_x} \omega_{v_x}}{k} - \beta + \frac{\beta}{k} \sum_{v_j \in EP'} \left(1 - \frac{d(v_j, v_x)}{D}\right) \\ &= \frac{\beta}{k} \sum_{v_i \in EP' - EP} \left(1 - \frac{d(v_i, v_x)}{D}\right) \end{aligned} \tag{2}$$

Since $\beta \in \mathbb{R}_{\geq 0}$, $k > 0$ and $d(v_i, v_x) \in [0, D]$, $\forall v_i, v_x \in V'$, $G(EP, v_x) - G(EP', v_x) \geq 0$ and hence $F$ is submodular. □

##### 1.2 Greedy algorithm to add SNPs to the set $EP$

The greedy algorithm below is used to add SNPs to set $EP$ (Nemhauser, 1978). This algorithm ensures a $\left(1 - \frac{1}{e}\right)$-factor approximation to the optimum solution when the submodular set function is also monotone non-decreasing and non-negative, These properties for $F$ are shown in Supplementary Text 1.1 and 1.2.

**Algorithm 1** Greedy Algorithm by Nemhauser, 1978

**Require:** Set function $F$, SNP set $V'$, desired subset size $k \leq |V'|$.
**Ensure:** Set $EP \subset V'$ such that $|EP| = k$.
1: $EP \leftarrow \emptyset$
2: **while** $|EP| < k$ **do**
3: $\quad EP \leftarrow EP \cup \underset{v_x \in V' \setminus EP}{\text{argmax}} F(EP \cup v_x)$
4: **end while**

##### 1.3 $F$ is non-negative and monotonically non-decreasing

To be able to use the greedy algorithm described in Supplementary Text 1.2 with the constant factor approximation guarantee, one should prove that the optimization function is monotone non-decreasing in addition to being submodular. The latter is shown in Supplementary Text 1.1. The former is proven here.

First, $F(EP) = 0$ if $EP = \emptyset$. Then, $F(EP)$ is a monotone non-decreasing and non-negative function if and only if the gain function $G$ (see Supplementary Text 1.1.) is always non-negative i.e. $G(EP, v_x) \geq 0$ for all sets $EP \subset V'$ and $v_x \in V' - EP$. Thus, the inequaity below should hold:

$$\frac{c_{v_x} \omega_{v_x}}{k} + \beta - \frac{\beta}{k} \sum_{v_i \in EP} \left(1 - \frac{d(v_i, v_x)}{D}\right) \geq 0 \tag{3}$$

Note that $c_{v_x} \geq 0$, $\omega_{v_x} \geq 1$, $\beta \geq 0$, $G(EP, v_x) \geq 0$. Then, this inequalty holds if $\beta \geq \frac{\beta}{k} \sum_{v_i \in EP} \left(1 - \frac{d(v_i, v_x)}{D}\right)$.The sum on the right hand side has a loose upper bound $|EP|$ which is when for every $v_i \in EP$, the distance between $v_i$ and $v_x$ is 0. As the inequality $k \geq |EP|$ holds, $G(EP, v_x) \geq 0$. $\square$

#### 2 Supplementary Figures

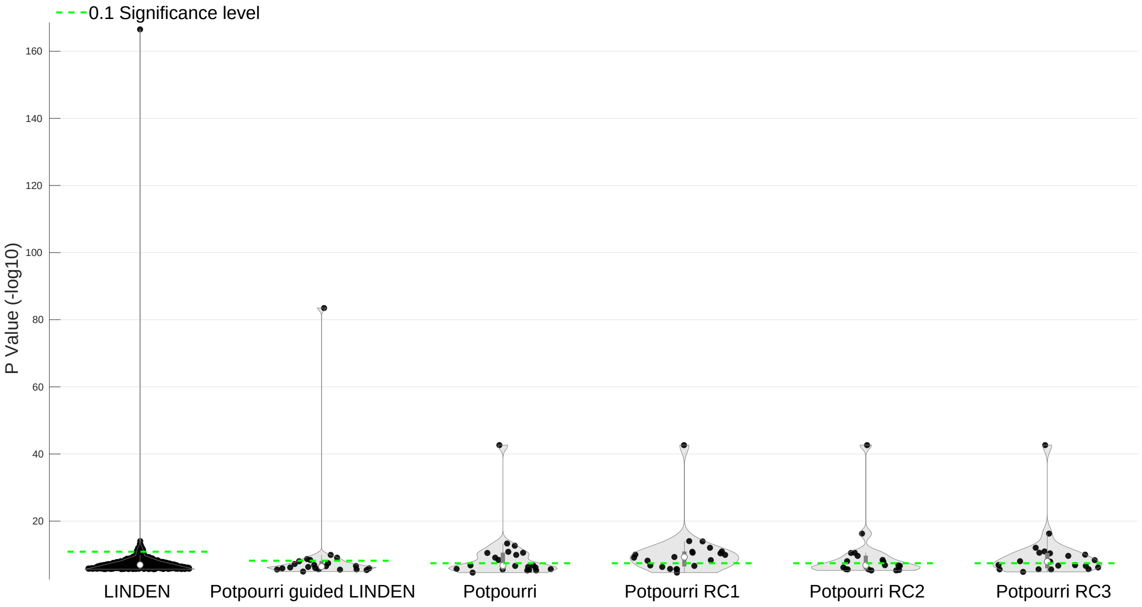


Figure 1: On the BD dataset, this figure compares the p-values of the selected SNP pairs by the following methods: (i) LINDEN, (ii) Potpourri *guided* LINDEN, (iii) Potpourri; and then also the variants of Potpourri which further promotes SNP pairs in regulatory and coding regions: (iv) Potpourri RC1, (v) Potpourri RC2, and finally (vi) Potpourri RC3. We show the significance levels (y-axis) of each reported pair (dots) given the Bonferroni-corrected significance threshold (0.1, green dashed lines). X − axis is just randomly assigned values to pairs for better visualization. Potpourri is run with $k = 750$ for all 5 related subplots. For RC1 and RC3 strategies $\omega$ is set to 1.31623, for Potpourri and RC2, it is set to 1. LINDEN is run with default parameters. For Potpourri *guided* LINDEN, tree formation is restricted to be done on distinct regions in Potpourri-selected regions **R** as described in Section **??**.

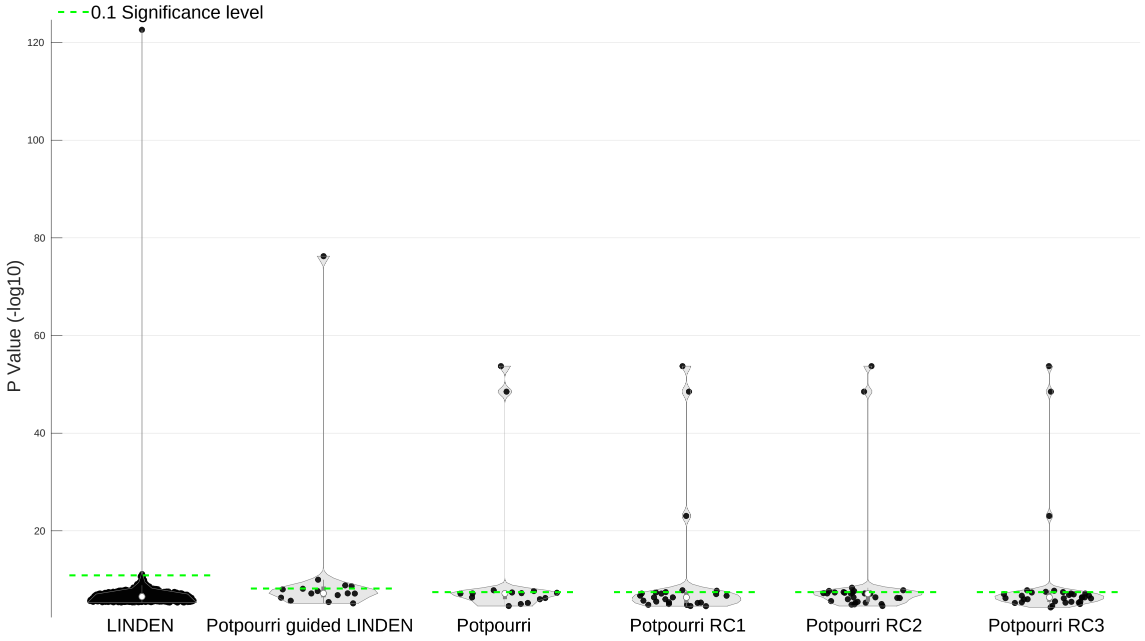


Figure 2: On the HT dataset, this figure compares the p-values of the selected SNP pairs by the following methods: (i) LINDEN, (ii) Potpourri *guided* LINDEN, (iii) Potpourri; and then also the variants of Potpourri which further promotes SNP pairs in regulatory and coding regions: (iv) Potpourri RC1, (v) Potpourri RC2, and finally (vi) Potpourri RC3. We show the significance levels (y-axis) of each reported pair (dots) given the Bonferroni-corrected significance threshold (0.1, green dashed lines). $X-$ axis is just randomly assigned values to pairs for better visualization. Potpourri is run with $k = 750$ for all 5 related subplots. For RC1 and RC3 strategies $\omega$ is set to 1.31623, for Potpourri and RC2, it is set to 1. LINDEN is run with default parameters. For Potpourri *guided* LINDEN, tree formation is restricted to be done on distinct regions in Potpourri-selected regions $\mathbf{R}$ as described in Section **??**.

### 3 Supplementary Tables

Table 1: Information of the T2D, BD and HT datasets which are used in our experiments.

| Dataset | # SNPs | # Cases | # Controls |
|---|---|---|---|
| **T2D** | 378016 | 1973 | 1498 |
| **HT** | 377456 | 1996 | 1498 |
| **BD** | 378008 | 1993 | 1498 |

Table 2: The number of genes in the Ensembl dataset and gene predictions dataset obtained from UCSC Genome Browser.

| | Gene Counts |
|---|---|
| Known protein-coding genes | 21370 |
| Novel protein-coding genes | 46 |
| Pseudogenes | 9899 |
| RNA genes | 5732 |
| Genscan gene predictions | 49796 |

Table 3: Results for BD dataset that compares LINDEN, Potpourri-guided LINDEN and Potpourri (with population co-cover). Number of pairs reported is the total number of reciprocally significant pairs returned by each method for varying number of selected SNPs. The number in parentheses denotes the significant pairs passing significance threshold (0.1) after Bonferroni correction based on the number of tests performed by each method. Bold denotes the best result for a given $k$ value. The table shows that the guidance of Potpourri substantially improves the precision of LINDEN. For all considered $k$ values, Potpourri provides the best precision values.

| **Method** | | **# Tested Loci** | **# Pairs Reported** | **Precision** |
|---|---|---|---|---|
| **LINDEN** | | 378008 | 885 (35) | 0.040 |
| **Potpourri guided LINDEN** | $k$ = *500* | 9387 | 30 (3) | 0.100 |
| | $k$ = *750* | 14098 | 22 (5) | 0.227 |
| | $k$ = *1000* | 17048 | 20 (7) | 0.350 |
| | $k$ = *1500* | 28161 | 47 (18) | 0.383 |
| | $k$ = *2000* | 36323 | 61 (13) | 0.213 |
| **Potpourri** | $k$ = *500* | 9382 | **33 (4)** | **0.121** |
| | $k$ = *750* | 14250 | **21 (9)** | **0.429** |
| | $k$ = *1000* | 17587 | **27 (15)** | **0.556** |
| | $k$ = *1500* | 28500 | **36 (22)** | **0.611** |
| | $k$ = *2000* | 36958 | **38 (17)** | **0.447** |

Table 4: Results for HT dataset that compares LINDEN, Potpourri-guided LINDEN and Potpourri (with population co-cover). Number of pairs reported is the total number of reciprocally significant pairs returned by each method for varying number of selected SNPs. The number in parentheses denotes the significant pairs passing significance threshold (0.1) after Bonferroni correction based on the number of tests performed by each method. Bold denotes the best result for a given $k$ value. The table shows that the guidance of Potpourri substantially improves the precision of LINDEN. For all but one considered $k$ values, Potpourri provides the best precision values.

| Method | | # Tested Loci | # Pairs Reported | Precision |
|---|---|---|---|---|
| **LINDEN** | | 377456 | 1022 (2) | 0.002 |
| **Potpourri guided LINDEN** | $k = 500$ | 8744 | 22 (6) | 0.272 |
| | $k = 750$ | 11886 | 15 (4) | 0.267 |
| | $k = 1000$ | 18953 | 31 (7) | 0.226 |
| | $k = 1500$ | 27275 | **40 (8)** | **0.200** |
| | $k = 2000$ | 37979 | 82 (3) | 0.037 |
| **Potpourri** | $k = 500$ | 8982 | **15 (5)** | **0.333** |
| | $k = 750$ | 13996 | **15 (4)** | **0.267** |
| | $k = 1000$ | 19000 | **29 (7)** | **0.241** |
| | $k = 1500$ | 21131 | 41 (6) | 0.146 |
| | $k = 2000$ | 37838 | **81 (7)** | **0.086** |

Table 5: LINDEN results for various $d$ values on T2D, BD and HT datasets. Number of pairs reported is the total number of reciprocally significant pairs returned by LINDEN. The number in parentheses denotes the significant pairs passing significance threshold (0.1) after Bonferroni correction based on the number of tests performed by LINDEN for each dataset. Table shows that making LINDEN more conservative does not improve the precision.

| | **d** | | | | | |
|---|---|---|---|---|---|---|
| | **0.45** | | **0.9** | | **0.99** | |
| **Dataset** | **# Pairs Reported** | **Precision** | **# Pairs Reported** | **Precision** | **# Pairs Reported** | **Precision** |
| **T2D** | 1786 (5) | 0.0028 | 1717 (3) | 0.0017 | 768 (5) | 0.0065 |
| **BD** | 885 (35) | 0.040 | 669 (8) | 0.0120 | 323 (9) | 0.0279 |
| **HT** | 1022 (2) | 0.002 | 683 (2) | 0.0029 | 328 (4) | 0.0122 |

Table 6: This table compares LINDEN and Potpourri guided LINDEN for $k = 750$ in terms of number of internal and leaf node tests conducted on T2D, BD and HT datasets. $d$ parameter is set to 0.45 in all settings. Results show that Potpourri guidance is able to reduce both the number of internal node tests and the number of leaf node tests substantially.

| Dataset | Method | # SNPs | # Trees tested | # Internal tests | # Leaf tests | Exhaustive test ratio |
|---|---|---|---|---|---|---|
| **T2D** | ***LINDEN*** | 378016 | 123390 | 5561643905 | 2067646712 | 0.107191 |
| | ***Potpourri guided LINDEN*** | 14146 | 5517 | 10223962 | 5459361 | 0.169076 |
| **BD** | ***LINDEN*** | 378008 | 123365 | 5560833906 | 2065965712 | 0.107561 |
| | ***Potpourri guided LINDEN*** | 14250 | 5513 | 10185791 | 5503138 | 0.169909 |
| **HT** | ***LINDEN*** | 377456 | 123213 | 5549612478 | 2057575718 | 0.107187 |
| | ***Potpourri guided LINDEN*** | 13996 | 5429 | 9983212 | 5257728 | 0.166377 |

Table 7: The table compares Potpourri results with three strategies (RC1, RC2,and RC3) to promote regulatory and coding regions on the T2D dataset. For RC1 and RC3 $\omega$ is set to 2. Number of pairs reported is the total number of reciprocally significant pairs. The number in parentheses denotes the significant pairs passing significance threshold (0.1) after Bonferroni correction based on the number of tests performed by each method. Bold denotes the best result for a given $k$ value.

| **Method** | | **# Tested Loci** | **# Reported (Rec. Sig.)** | **Precision** |
|---|---|---|---|---|
| **Potpourri** | $k$ = *500* | 9250 | **12 (8)** | **0.667** |
| | $k$ = *750* | 14145 | **23 (15)** | **0.652** |
| | $k$ = *1000* | 18943 | 31 (20) | 0.645 |
| **Potpourri RC1** | $k$ = *500* | 9362 | 13 (8) | 0.615 |
| | $k$ = *750* | 14250 | 28 (15) | 0.536 |
| | $k$ = *1000* | 18834 | 30 (14) | 0.467 |
| **Potpourri RC2** | $k$ = *500* | 9250 | 13 (7) | 0.538 |
| | $k$ = *750* | 14145 | 24 (14) | 0.583 |
| | $k$ = *1000* | 18943 | **33 (23)** | **0.697** |
| **Potpourri RC3** | $k$ = *500* | 9362 | 18 (7) | 0.389 |
| | $k$ = *750* | 14250 | 25 (13) | 0.520 |
| | $k$ = *1000* | 18834 | 31 (15) | 0.484 |

Table 8: The table compares Potpourri results with three strategies (RC1, RC2,and RC3) to promote regulatory and coding regions on the T2D dataset. For RC1 and RC3 $\omega$ is set to 1.56234. Number of pairs reported is the total number of reciprocally significant pairs. The number in parentheses denotes the significant pairs passing significance threshold (0.1) after Bonferroni correction based on the number of tests performed by each method. Bold denotes the best result for a given $k$ value.

| **Method** | | **# Tested Loci** | **# Reported (Rec. Sig.)** | **Precision** |
|---|---|---|---|---|
| **Potpourri** | $k$ = *500* | 9250 | **12 (8)** | **0.667** |
| | $k$ = *750* | 14145 | 23 (15) | 0.652 |
| | $k$ = *1000* | 18943 | 31 (20) | 0.645 |
| **Potpourri RC1** | $k$ = *500* | 9365 | 14 (9) | 0.643 |
| | $k$ = *750* | 14185 | **19 (14)** | **0.737** |
| | $k$ = *1000* | 18677 | 32 (17) | 0.531 |
| **Potpourri RC2** | $k$ = *500* | 9250 | 13 (7) | 0.538 |
| | $k$ = *750* | 14145 | 24 (14) | 0.583 |
| | $k$ = *1000* | 18943 | **33 (23)** | **0.697** |
| **Potpourri RC3** | $k$ = *500* | 9365 | 15 (6) | 0.400 |
| | $k$ = *750* | 14185 | 20 (13) | 0.650 |
| | $k$ = *1000* | 18677 | 32 (17) | 0.531 |

Table 9: The table compares Potpourri results with three strategies (RC1, RC2,and RC3) to promote regulatory and coding regions on the BD dataset. For RC1 and RC3 $\omega$ is set to 2. Number of pairs reported is the total number of reciprocally significant pairs. The number in parentheses denotes the significant pairs passing significance threshold (0.1) after Bonferroni correction based on the number of tests performed by each method. Bold denotes the best result for a given $k$ value.

| **Method** | | **# Tested Loci** | **# Reported (Rec. Sig.)** | **Precision** |
|---|---|---|---|---|
| **Potpourri** | $k$ = *500* | 9382 | 33 (4) | 0.121 |
| | $k$ = *750* | 14250 | 21 (9) | 0.429 |
| | $k$ = *1000* | 17587 | 27 (15) | 0.556 |
| **Potpourri RC1** | $k$ = *500* | 9262 | **16 (4)** | **0.250** |
| | $k$ = *750* | 14199 | **24 (11)** | **0.458** |
| | $k$ = *1000* | 18660 | 22 (13) | 0.591 |
| **Potpourri RC2** | $k$ = *500* | 9382 | 35 (4) | 0.114 |
| | $k$ = *750* | 14250 | 18 (8) | 0.444 |
| | $k$ = *1000* | 17587 | **23 (15)** | **0.652** |
| **Potpourri RC3** | $k$ = *500* | 9262 | 14 (1) | 0.071 |
| | $k$ = *750* | 14199 | 19 (8) | 0.421 |
| | $k$ = *1000* | 18660 | 23 (11) | 0.478 |

Table 10: The table compares Potpourri results with three strategies (RC1, RC2,and RC3) to promote regulatory and coding regions on the BD dataset. For RC1 and RC3 $\omega$ is set to 1.56234. Number of pairs reported is the total number of reciprocally significant pairs. The number in parentheses denotes the significant pairs passing significance threshold (0.1) after Bonferroni correction based on the number of tests performed by each method. Bold denotes the best result for a given $k$ value.

| **Method** | | **# Tested Loci** | **# Reported (Rec. Sig.)** | **Precision** |
|---|---|---|---|---|
| **Potpourri** | $k$ = *500* | 9382 | 33 (4) | 0.121 |
| | $k$ = *750* | 14250 | 21 (9) | 0.429 |
| | $k$ = *1000* | 17587 | 27 (15) | 0.556 |
| **Potpourri RC1** | $k$ = *500* | 9325 | **17 (3)** | **0.176** |
| | $k$ = *750* | 13932 | 14 (6) | 0.429 |
| | $k$ = *1000* | 16800 | 23 (14) | 0.609 |
| **Potpourri RC2** | $k$ = *500* | 9382 | 35 (4) | 0.114 |
| | $k$ = *750* | 14250 | **18 (8)** | **0.444** |
| | $k$ = *1000* | 17587 | **23 (15)** | **0.652** |
| **Potpourri RC3** | $k$ = *500* | 9325 | 18 (3) | 0.167 |
| | $k$ = *750* | 13932 | 17 (7) | 0.412 |
| | $k$ = *1000* | 16800 | 21 (11) | 0.524 |

Table 11: The table compares Potpourri results with three strategies (RC1, RC2,and RC3) to promote regulatory and coding regions on the BD dataset. For RC1 and RC3 $\omega$ is set to 1.31623. Number of pairs reported is the total number of reciprocally significant pairs. The number in parentheses denotes the significant pairs passing significance threshold (0.1) after Bonferroni correction based on the number of tests performed by each method. Bold denotes the best result for a given $k$ value.

| **Method** | | **# Tested Loci** | **# Reported (Rec. Sig.)** | **Precision** |
|---|---|---|---|---|
| **Potpourri** | $k$ = *500* | 9382 | 33 (4) | 0.121 |
| | $k$ = *750* | 14250 | 21 (9) | 0.429 |
| | $k$ = *1000* | 17587 | 27 (15) | 0.556 |
| **Potpourri RC1** | $k$ = *500* | 9304 | **18 (4)** | **0.222** |
| | $k$ = *750* | 14250 | **21 (14)** | **0.667** |
| | $k$ = *1000* | 16828 | 24 (14) | 0.583 |
| **Potpourri RC2** | $k$ = *500* | 9382 | 35 (4) | 0.114 |
| | $k$ = *750* | 14250 | 18 (8) | 0.444 |
| | $k$ = *1000* | 17587 | **23 (15)** | **0.652** |
| **Potpourri RC3** | $k$ = *500* | 9304 | 18 (2) | 0.111 |
| | $k$ = *750* | 14250 | 21 (11) | 0.524 |
| | $k$ = *1000* | 16828 | 24 (14) | 0.583 |

Table 12: The table compares Potpourri results with three strategies (RC1, RC2,and RC3) to promote regulatory and coding regions on the HT dataset. For RC1 and RC3 $\omega$ is set to 2. Number of pairs reported is the total number of reciprocally significant pairs. The number in parentheses denotes the significant pairs passing significance threshold (0.1) after Bonferroni correction based on the number of tests performed by each method. Bold denotes the best result for a given $k$ value.

| **Method** | | **# Tested Loci** | **# Reported (Rec. Sig.)** | **Precision** |
|---|---|---|---|---|
| **Potpourri** | $k$ = *500* | 8982 | 15 (5) | 0.333 |
| | $k$ = *750* | 13996 | 15 (4) | 0.267 |
| | $k$ = *1000* | 19000 | **29 (7)** | **0.241** |
| **Potpourri RC1** | $k$ = *500* | 9452 | 22 (4) | 0.182 |
| | $k$ = *750* | 12454 | 22 (4) | 0.182 |
| | $k$ = *1000* | 19000 | 37 (7) | 0.189 |
| **Potpourri RC2** | $k$ = *500* | 8982 | **20 (7)** | **0.350** |
| | $k$ = *750* | 13996 | 28 (7) | 0.250 |
| | $k$ = *1000* | 19000 | 45 (7) | 0.156 |
| **Potpourri RC3** | $k$ = *500* | 9452 | 26 (6) | 0.231 |
| | $k$ = *750* | 12454 | **40 (11)** | **0.275** |
| | $k$ = *1000* | 19000 | 48 (8) | 0.167 |

Table 13: The table compares Potpourri results with three strategies (RC1, RC2,and RC3) to promote regulatory and coding regions on the HT dataset. For RC1 and RC3 $\omega$ is set to 1.56234. Number of pairs reported is the total number of reciprocally significant pairs. The number in parentheses denotes the significant pairs passing significance threshold (0.1) after Bonferroni correction based on the number of tests performed by each method. Bold denotes the best result for a given $k$ value.

| **Method** | | **# Tested Loci** | **# Reported (Rec. Sig.)** | **Precision** |
|---|---|---|---|---|
| **Potpourri** | $k$ = *500* | 8982 | 15 (5) | 0.333 |
| | $k$ = *750* | 13996 | **15 (4)** | **0.267** |
| | $k$ = *1000* | 19000 | **29 (7)** | **0.241** |
| **Potpourri RC1** | $k$ = *500* | 9379 | 15 (3) | 0.200 |
| | $k$ = *750* | 11573 | 21 (4) | 0.190 |
| | $k$ = *1000* | 18935 | 31 (6) | 0.194 |
| **Potpourri RC2** | $k$ = *500* | 8982 | **20 (7)** | **0.350** |
| | $k$ = *750* | 13996 | 28 (7) | 0.250 |
| | $k$ = *1000* | 19000 | 45 (7) | 0.156 |
| **Potpourri RC3** | $k$ = *500* | 9379 | 23 (7) | 0.304 |
| | $k$ = *750* | 11573 | 34 (8) | 0.235 |
| | $k$ = *1000* | 18935 | 47 (7) | 0.149 |

Table 14: The table compares Potpourri results with three strategies (RC1, RC2,and RC3) to promote regulatory and coding regions on the HT dataset. For RC1 and RC3 $\omega$ is set to 1.31623. Number of pairs reported is the total number of reciprocally significant pairs. The number in parentheses denotes the significant pairs passing significance threshold (0.1) after Bonferroni correction based on the number of tests performed by each method. Bold denotes the best result for a given $k$ value.

| **Method** | | **# Tested Loci** | **# Reported (Rec. Sig.)** | **Precision** |
|---|---|---|---|---|
| **Potpourri** | $k$ = *500* | 8982 | 15 (5) | 0.333 |
| | $k$ = *750* | 13996 | **15 (4)** | **0.267** |
| | $k$ = *1000* | 19000 | 29 (7) | 0.241 |
| **Potpourri RC1** | $k$ = *500* | 8557 | 12 (2) | 0.167 |
| | $k$ = *750* | 14202 | 25 (6) | 0.240 |
| | $k$ = *1000* | 18509 | **23 (6)** | **0.261** |
| **Potpourri RC2** | $k$ = *500* | 8982 | **20 (7)** | **0.350** |
| | $k$ = *750* | 13996 | 28 (7) | 0.250 |
| | $k$ = *1000* | 19000 | 45 (7) | 0.156 |
| **Potpourri RC3** | $k$ = *500* | 8557 | 26 (8) | 0.308 |
| | $k$ = *750* | 14202 | 33 (7) | 0.212 |
| | $k$ = *1000* | 18509 | 35 (6) | 0.171 |

Table 15: For T2D dataset, reciprocally significant SNP pairs found by Potpourri - $k = 750$ and multiple hypothesis testing threshold is $\sim 3.56E - 08$

| **Chi-Sq.** | **P-value** | **SNP1 Chr** | **SNP2 Chr** | **SNP1 - Pos** | **SNP2 - Pos** | **SNP1 - rsID** | **SNP2 - rsID** |
|---|---|---|---|---|---|---|---|
| 188.321 | 8.89E-37 | 4 | 4 | 44958383 | 86618019 | rs28576190 | rs41401555 |
| 152.887 | 2.35E-29 | 1 | 4 | 156523434 | 177984277 | rs41362848 | rs41497848 |
| 71.2098 | 1.30E-12 | 10 | 11 | 71976274 | 26799868 | rs10823561 | rs12363428 |
| 70.9198 | 1.48E-12 | 1 | 3 | 180643701 | 114040644 | rs10797774 | rs7642984 |
| 69.4477 | 2.90E-12 | 10 | 16 | 82112773 | 9970587 | rs1538818 | rs4780746 |
| 68.0659 | 5.45E-12 | 17 | 22 | 13840981 | 36004493 | rs9897978 | rs12170152 |
| 63.2266 | 4.91E-11 | 2 | 13 | 193168479 | 42909766 | rs10497732 | rs17635556 |
| 59.8476 | 2.25E-10 | 4 | 10 | 20884293 | 125911953 | rs2322884 | rs11245116 |
| 59.2771 | 2.91E-10 | 8 | 10 | 77507371 | 114771287 | rs12548378 | rs10787472 |
| 58.5839 | 3.98E-10 | 8 | 9 | 25108289 | 86674433 | rs6996838 | rs4877289 |
| 57.113 | 7.69E-10 | 1 | 2 | 89495932 | 606686 | rs2209307 | rs7571872 |
| 51.8099 | 8.14E-09 | 13 | 22 | 86179358 | 47701798 | rs3015528 | rs8141372 |
| 51.4831 | 9.40E-09 | 1 | 10 | 183726314 | 73442342 | rs1039561 | rs731027 |
| 50.6357 | 1.37E-08 | 4 | 5 | 189725728 | 160929289 | rs1600203 | rs1444741 |
| 50.0637 | 1.76E-08 | 8 | 23 | 5759880 | 39675267 | rs2840450 | rs6520600 |
| 46.9504 | 6.88E-08 | 1 | 5 | 69745492 | 170333431 | rs17395543 | rs4868053 |
| 46.7405 | 7.54E-08 | 20 | 21 | 37776057 | 43194058 | rs292877 | rs8131547 |
| 45.7535 | 1.16E-07 | 15 | 20 | 31985488 | 12098580 | rs572917 | rs13044874 |
| 43.6802 | 2.84E-07 | 5 | 17 | 815435 | 12472981 | rs429739 | rs11655394 |
| 43.4085 | 3.19E-07 | 4 | 11 | 56621813 | 109803757 | rs2412720 | rs4753891 |
| 39.5984 | 1.63E-06 | 3 | 8 | 32338002 | 27025518 | rs9833771 | rs4621816 |
| 35.518 | 9.05E-06 | 2 | 9 | 221798041 | 95407641 | rs17349283 | rs2994370 |
| 32.7495 | 2.83E-05 | 12 | 21 | 113031202 | 43074023 | rs2195220 | rs9976865 |

Table 16: For T2D dataset, reciprocally significant SNP pairs found by Potpourri RC1 - $k = 750$ and $\omega = 1.31623$ and multiple hypothesis testing threshold is $\sim 3.57E - 08$

| **Chi-Sq.** | **P-value** | **SNP1 Chr** | **SNP2 Chr** | **SNP1 - Pos** | **SNP2 - Pos** | **SNP1 - rsID** | **SNP2 - rsID** |
|---|---|---|---|---|---|---|---|
| 188.321 | 8.89E-37 | 4 | 4 | 44958383 | 86618019 | rs28576190 | rs41401555 |
| 152.887 | 2.35E-29 | 1 | 4 | 156523434 | 177984277 | rs41362848 | rs41497848 |
| 86.4075 | 1.16E-15 | 5 | 17 | 108928074 | 76399520 | rs4504431 | rs2333990 |
| 70.9198 | 1.48E-12 | 1 | 3 | 180643701 | 114040644 | rs10797774 | rs7642984 |
| 69.6512 | 2.64E-12 | 10 | 14 | 100181866 | 63478768 | rs11599112 | rs9323443 |
| 69.4477 | 2.90E-12 | 10 | 16 | 82112773 | 9970587 | rs1538818 | rs4780746 |
| 61.3099 | 1.17E-10 | 11 | 15 | 95466766 | 72402560 | rs4753717 | rs11072478 |
| 59.8476 | 2.25E-10 | 4 | 10 | 20884293 | 125911953 | rs2322884 | rs11245116 |
| 58.5839 | 3.98E-10 | 8 | 9 | 25108289 | 86674433 | rs6996838 | rs4877289 |
| 51.4831 | 9.40E-09 | 1 | 10 | 183726314 | 73442342 | rs1039561 | rs731027 |
| 50.0637 | 1.76E-08 | 8 | 23 | 5759880 | 39675267 | rs2840450 | rs6520600 |
| 49.5518 | 2.20E-08 | 9 | 13 | 36912960 | 111039550 | rs2297109 | rs4773395 |
| 46.9504 | 6.88E-08 | 1 | 5 | 69745492 | 170333431 | rs17395543 | rs4868053 |
| 46.7405 | 7.54E-08 | 20 | 21 | 37776057 | 43194058 | rs292877 | rs8131547 |
| 35.518 | 9.05E-06 | 2 | 9 | 221798041 | 95407641 | rs17349283 | rs2994370 |

Table 17: For T2D dataset, reciprocally significant SNP pairs found by Potpourri RC2 - $k = 750$ and multiple hypothesis testing threshold is $\sim 3.56E - 08$

| Chi-Sq. | P-value | SNP1 Chr | SNP2 Chr | SNP1 - Pos | SNP2 - Pos | SNP1 - rsID | SNP2 - rsID |
|---|---|---|---|---|---|---|---|
| 188.321 | 8.89E-37 | 4 | 4 | 44958383 | 86618019 | rs28576190 | rs41401555 |
| 71.2098 | 1.30E-12 | 10 | 11 | 71976274 | 26799868 | rs10823561 | rs12363428 |
| 70.9198 | 1.48E-12 | 1 | 3 | 180643701 | 114040644 | rs10797774 | rs7642984 |
| 69.4477 | 2.90E-12 | 10 | 16 | 82112773 | 9970587 | rs1538818 | rs4780746 |
| 68.0659 | 5.45E-12 | 17 | 22 | 13840981 | 36004493 | rs9897978 | rs12170152 |
| 63.2266 | 4.91E-11 | 2 | 13 | 193168479 | 42909766 | rs10497732 | rs17635556 |
| 59.8476 | 2.25E-10 | 4 | 10 | 20884293 | 125911953 | rs2322884 | rs11245116 |
| 59.2771 | 2.91E-10 | 8 | 10 | 77507371 | 114771287 | rs12548378 | rs10787472 |
| 57.113 | 7.69E-10 | 1 | 2 | 89495932 | 606686 | rs2209307 | rs7571872 |
| 53.4052 | 4.02E-09 | 5 | 23 | 153621510 | 148374123 | rs11167666 | rs5936294 |
| 51.8099 | 8.14E-09 | 13 | 22 | 86179358 | 47701798 | rs3015528 | rs8141372 |
| 51.4831 | 9.40E-09 | 1 | 10 | 183726314 | 73442342 | rs1039561 | rs731027 |
| 50.6357 | 1.37E-08 | 4 | 5 | 189725728 | 160929289 | rs1600203 | rs1444741 |
| 50.0637 | 1.76E-08 | 8 | 23 | 5759880 | 39675267 | rs2840450 | rs6520600 |
| 46.7405 | 7.54E-08 | 20 | 21 | 37776057 | 43194058 | rs292877 | rs8131547 |
| 45.852 | 1.11E-07 | 11 | 16 | 125282266 | 76766570 | rs485878 | rs11643308 |
| 45.7535 | 1.16E-07 | 15 | 20 | 31985488 | 12098580 | rs572917 | rs13044874 |
| 43.6802 | 2.84E-07 | 5 | 17 | 815435 | 12472981 | rs429739 | rs11655394 |
| 41.154 | 8.40E-07 | 2 | 10 | 173065773 | 87533919 | rs7564591 | rs2949386 |
| 38.8487 | 2.24E-06 | 4 | 5 | 132765095 | 143234501 | rs350965 | rs10075045 |
| 35.518 | 9.05E-06 | 2 | 9 | 221798041 | 95407641 | rs17349283 | rs2994370 |
| 34.0265 | 1.68E-05 | 12 | 14 | 2848515 | 36558759 | rs2074985 | rs9806037 |
| 33.8518 | 1.80E-05 | 4 | 6 | 56577074 | 26124441 | rs2611826 | rs199750 |
| 32.7495 | 2.83E-05 | 12 | 21 | 113031202 | 43074023 | rs2195220 | rs9976865 |

Table 18: For T2D dataset, reciprocally significant SNP pairs found by Potpourri RC3 - $k = 750$ and $\omega = 1.31623$ and multiple hypothesis testing threshold is $\sim 3.57E - 08$

| Chi-Sq. | P-value | SNP1 Chr | SNP2 Chr | SNP1 - Pos | SNP2 - Pos | SNP1 - rsID | SNP2 - rsID |
|---|---|---|---|---|---|---|---|
| 188.321 | 8.89E-37 | 4 | 4 | 44958383 | 86618019 | rs28576190 | rs41401555 |
| 70.9198 | 1.48E-12 | 1 | 3 | 180643701 | 114040644 | rs10797774 | rs7642984 |
| 69.6512 | 2.64E-12 | 10 | 14 | 100181866 | 63478768 | rs11599112 | rs9323443 |
| 69.4477 | 2.90E-12 | 10 | 16 | 82112773 | 9970587 | rs1538818 | rs4780746 |
| 61.3099 | 1.17E-10 | 11 | 15 | 95466766 | 72402560 | rs4753717 | rs11072478 |
| 59.8476 | 2.25E-10 | 4 | 10 | 20884293 | 125911953 | rs2322884 | rs11245116 |
| 57.9618 | 5.26E-10 | 8 | 23 | 2301019 | 43874622 | rs7827690 | rs5906691 |
| 53.4052 | 4.02E-09 | 5 | 23 | 153621510 | 148374123 | rs11167666 | rs5936294 |
| 51.4831 | 9.40E-09 | 1 | 10 | 183726314 | 73442342 | rs1039561 | rs731027 |
| 50.0637 | 1.76E-08 | 8 | 23 | 5759880 | 39675267 | rs2840450 | rs6520600 |
| 49.5518 | 2.20E-08 | 9 | 13 | 36912960 | 111039550 | rs2297109 | rs4773395 |
| 46.7405 | 7.54E-08 | 20 | 21 | 37776057 | 43194058 | rs292877 | rs8131547 |
| 41.154 | 8.40E-07 | 2 | 10 | 173065773 | 87533919 | rs7564591 | rs2949386 |
| 35.518 | 9.05E-06 | 2 | 9 | 221798041 | 95407641 | rs17349283 | rs2994370 |
| 34.0265 | 1.68E-05 | 12 | 14 | 2848515 | 36558759 | rs2074985 | rs9806037 |

Table 19: For BD dataset, reciprocally significant SNP pairs found by Potpourri - $k = 750$ and multiple hypothesis testing threshold is $\sim 3.56E - 08$

| Chi-Sq. | P-value | SNP1 Chr | SNP2 Chr | SNP1 - Pos | SNP2 - Pos | SNP1 - rsID | SNP2 - rsID |
|---|---|---|---|---|---|---|---|
| 219.583 | 2.29E-43 | 2 | 17 | 191990530 | 65201507 | rs41464947 | rs41432448 |
| 78.4245 | 4.69E-14 | 1 | 11 | 232548301 | 59134119 | rs3001707 | rs11603417 |
| 75.1371 | 2.14E-13 | 9 | 13 | 34078701 | 107336856 | rs11793082 | rs1509095 |
| 65.9714 | 1.41E-11 | 11 | 16 | 6262553 | 2439012 | rs11040843 | rs8060813 |
| 64.586 | 2.65E-11 | 3 | 17 | 196184043 | 31427052 | rs849891 | rs862807 |
| 64.2557 | 3.08E-11 | 6 | 10 | 91726062 | 119194975 | rs7775358 | rs2768314 |
| 61.3257 | 1.16E-10 | 10 | 12 | 80683764 | 46662480 | rs1250604 | rs12811832 |
| 57.0785 | 7.81E-10 | 2 | 14 | 86578578 | 20795494 | rs6725080 | rs17197037 |
| 53.5585 | 3.75E-09 | 2 | 18 | 11954943 | 2292770 | rs4027132 | rs7240747 |
| 45.1211 | 1.52E-07 | 15 | 23 | 89872962 | 7980438 | rs4932551 | rs5934353 |
| 44.9839 | 1.62E-07 | 1 | 3 | 58899674 | 7177533 | rs7523134 | rs1818033 |
| 44.2482 | 2.22E-07 | 5 | 13 | 672022 | 89239100 | rs1617985 | rs9522711 |
| 42.5742 | 4.57E-07 | 2 | 4 | 37022553 | 71014262 | rs10193295 | rs6856412 |
| 41.7297 | 6.57E-07 | 9 | 23 | 36979054 | 138898572 | rs4880048 | rs5953923 |
| 39.4318 | 1.75E-06 | 9 | 17 | 116362363 | 76396399 | rs10817638 | rs2672887 |
| 39.0046 | 2.10E-06 | 3 | 14 | 172586501 | 45407498 | rs902956 | rs444248 |
| 38.6986 | 2.39E-06 | 3 | 3 | 26750767 | 121419173 | rs1386886 | rs6778180 |
| 37.6767 | 3.67E-06 | 4 | 11 | 183409195 | 134377514 | rs11724819 | rs7102094 |
| 36.9293 | 5.02E-06 | 4 | 7 | 122883773 | 87040551 | rs28532673 | rs10264990 |
| 36.9117 | 5.06E-06 | 6 | 12 | 72226595 | 81094156 | rs9351813 | rs11115212 |
| 33.3186 | 2.24E-05 | 11 | 23 | 37103169 | 12507198 | rs7127861 | rs12012548 |

Table 20: For BD dataset, reciprocally significant SNP pairs found by Potpourri RC1 - $k = 750$ and $\omega = 1.31623$ and multiple hypothesis testing threshold is $\sim 3.56E - 08$.

| Chi-Sq. | P-value | SNP1 Chr | SNP2 Chr | SNP1 - Pos | SNP2 - Pos | SNP1 - rsID | SNP2 - rsID |
|---|---|---|---|---|---|---|---|
| 219.583 | 2.29E-43 | 2 | 17 | 191990530 | 65201507 | rs41464947 | rs41432448 |
| 81.9403 | 9.23E-15 | 11 | 20 | 59134119 | 15396092 | rs11603417 | rs1884706 |
| 81.5692 | 1.10E-14 | 5 | 17 | 53508615 | 57095575 | rs277336 | rs7220740 |
| 72.0609 | 8.77E-13 | 1 | 18 | 163690500 | 11187112 | rs7536062 | rs12962470 |
| 66.547 | 1.09E-11 | 2 | 7 | 172823460 | 5358228 | rs4435421 | rs3801048 |
| 65.9714 | 1.41E-11 | 11 | 16 | 6262553 | 2439012 | rs11040843 | rs8060813 |
| 64.586 | 2.65E-11 | 3 | 17 | 196184043 | 31427052 | rs849891 | rs862807 |
| 63.6721 | 4.01E-11 | 2 | 3 | 80504972 | 63565854 | rs7557554 | rs1876092 |
| 61.6663 | 9.94E-11 | 9 | 9 | 21913279 | 27723456 | rs10811638 | rs1857080 |
| 61.3257 | 1.16E-10 | 10 | 12 | 80683764 | 46662480 | rs1250604 | rs12811832 |
| 58.084 | 4.98E-10 | 7 | 15 | 5308096 | 81153313 | rs4299914 | rs8043401 |
| 57.0785 | 7.81E-10 | 2 | 14 | 86578578 | 20795494 | rs6725080 | rs17197037 |
| 53.2643 | 4.27E-09 | 1 | 4 | 83092765 | 15607568 | rs12407742 | rs6449211 |
| 52.4183 | 6.22E-09 | 3 | 9 | 134399302 | 135448225 | rs6439389 | rs3025299 |
| 44.9839 | 1.62E-07 | 1 | 3 | 58899674 | 7177533 | rs7523134 | rs1818033 |
| 44.2482 | 2.22E-07 | 5 | 13 | 672022 | 89239100 | rs1617985 | rs9522711 |
| 42.5742 | 4.57E-07 | 2 | 4 | 37022553 | 71014262 | rs10193295 | rs6856412 |
| 39.4318 | 1.75E-06 | 9 | 17 | 116362363 | 76396399 | rs10817638 | rs2672887 |
| 39.0046 | 2.10E-06 | 3 | 14 | 172586501 | 45407498 | rs902956 | rs444248 |
| 38.6986 | 2.39E-06 | 3 | 3 | 26750767 | 121419173 | rs1386886 | rs6778180 |
| 33.3186 | 2.24E-05 | 11 | 23 | 37103169 | 12507198 | rs7127861 | rs12012548 |

Table 21: For BD dataset, reciprocally significant SNP pairs found by Potpourri RC2 - $k = 750$ and and multiple hypothesis testing threshold is $\sim 3.56E-08$

| **Chi-Sq.** | **P-value** | **SNP1 Chr** | **SNP2 Chr** | **SNP1 - Pos** | **SNP2 - Pos** | **SNP1 - rsID** | **SNP2 - rsID** |
|---|---|---|---|---|---|---|---|
| 219.583 | 2.29E-43 | 2 | 17 | 191990530 | 65201507 | rs41464947 | rs41432448 |
| 93.023 | 5.29E-17 | 16 | 23 | 47086325 | 140870141 | rs41334049 | rs11795560 |
| 64.586 | 2.65E-11 | 3 | 17 | 196184043 | 31427052 | rs849891 | rs862807 |
| 64.2557 | 3.08E-11 | 6 | 10 | 91726062 | 119194975 | rs7775358 | rs2768314 |
| 59.7868 | 2.32E-10 | 10 | 20 | 7441884 | 24363277 | rs638145 | rs6049681 |
| 53.5585 | 3.75E-09 | 2 | 18 | 11954943 | 2292770 | rs4027132 | rs7240747 |
| 51.5925 | 8.96E-09 | 2 | 23 | 235155540 | 150611004 | rs1526107 | rs5925006 |
| 51.0117 | 1.16E-08 | 7 | 15 | 125067319 | 31377909 | rs7782495 | rs8031347 |
| 45.4226 | 1.34E-07 | 8 | 23 | 26416283 | 67899848 | rs13258081 | rs792952 |
| 45.1211 | 1.52E-07 | 15 | 23 | 89872962 | 7980438 | rs4932551 | rs5934353 |
| 44.2482 | 2.22E-07 | 5 | 13 | 672022 | 89239100 | rs1617985 | rs9522711 |
| 41.7959 | 6.39E-07 | 5 | 6 | 91479145 | 5458549 | rs10942660 | rs12204218 |
| 39.0046 | 2.10E-06 | 3 | 14 | 172586501 | 45407498 | rs902956 | rs444248 |
| 38.8624 | 2.23E-06 | 2 | 14 | 37086769 | 73267706 | rs17038861 | rs7140655 |
| 38.6986 | 2.39E-06 | 3 | 3 | 26750767 | 121419173 | rs1386886 | rs6778180 |
| 37.6767 | 3.67E-06 | 4 | 11 | 183409195 | 134377514 | rs11724819 | rs7102094 |
| 37.0117 | 4.85E-06 | 11 | 18 | 115696852 | 44489439 | rs4938261 | rs1384226 |
| 36.9117 | 5.06E-06 | 6 | 12 | 72226595 | 81094156 | rs9351813 | rs11115212 |

Table 22: For BD dataset, reciprocally significant SNP pairs found by Potpourri RC3 - $k = 750$ and $\omega = 1.31623$ and multiple hypothesis testing threshold is $\sim 3.56E-08$

| **Chi-Sq.** | **P-value** | **SNP1 Chr** | **SNP2 Chr** | **SNP1 - Pos** | **SNP2 - Pos** | **SNP1 - rsID** | **SNP2 - rsID** |
|---|---|---|---|---|---|---|---|
| 219.583 | 2.29E-43 | 2 | 17 | 191990530 | 65201507 | rs41464947 | rs41432448 |
| 93.023 | 5.29E-17 | 16 | 23 | 47086325 | 140870141 | rs41334049 | rs11795560 |
| 72.0609 | 8.77E-13 | 1 | 18 | 163690500 | 11187112 | rs7536062 | rs12962470 |
| 66.547 | 1.09E-11 | 2 | 7 | 172823460 | 5358228 | rs4435421 | rs3801048 |
| 64.586 | 2.65E-11 | 3 | 17 | 196184043 | 31427052 | rs849891 | rs862807 |
| 63.6721 | 4.01E-11 | 2 | 3 | 80504972 | 63565854 | rs7557554 | rs1876092 |
| 61.6663 | 9.94E-11 | 9 | 9 | 21913279 | 27723456 | rs10811638 | rs1857080 |
| 59.7868 | 2.32E-10 | 10 | 20 | 7441884 | 24363277 | rs638145 | rs6049681 |
| 53.2643 | 4.27E-09 | 1 | 4 | 83092765 | 15607568 | rs12407742 | rs6449211 |
| 51.5925 | 8.96E-09 | 2 | 23 | 235155540 | 150611004 | rs1526107 | rs5925006 |
| 51.0117 | 1.16E-08 | 7 | 15 | 125067319 | 31377909 | rs7782495 | rs8031347 |
| 45.7479 | 1.16E-07 | 11 | 23 | 95048677 | 18757736 | rs7932590 | rs181150 |
| 45.4226 | 1.34E-07 | 8 | 23 | 26416283 | 67899848 | rs13258081 | rs792952 |
| 44.6958 | 1.83E-07 | 7 | 23 | 42529578 | 30435432 | rs10260633 | rs5972177 |
| 44.2482 | 2.22E-07 | 5 | 13 | 672022 | 89239100 | rs1617985 | rs9522711 |
| 41.7959 | 6.39E-07 | 5 | 6 | 91479145 | 5458549 | rs10942660 | rs12204218 |
| 39.0046 | 2.10E-06 | 3 | 14 | 172586501 | 45407498 | rs902956 | rs444248 |
| 38.8624 | 2.23E-06 | 2 | 14 | 37086769 | 73267706 | rs17038861 | rs7140655 |
| 38.6986 | 2.39E-06 | 3 | 3 | 26750767 | 121419173 | rs1386886 | rs6778180 |
| 38.6336 | 2.45E-06 | 12 | 18 | 130242349 | 6867710 | rs4644706 | rs547779 |
| 34.5439 | 1.35E-05 | 6 | 11 | 76683476 | 78664406 | rs6914716 | rs509249 |

Table 23: For HT dataset, reciprocally significant SNP pairs found by Potpourri - $k = 750$ and multiple hypothesis testing threshold is $\sim 3.56E - 08$

| **Chi-Sq.** | **P-value** | **SNP1 Chr** | **SNP2 Chr** | **SNP1 - Pos** | **SNP2 - Pos** | **SNP1 - rsID** | **SNP2 - rsID** |
|---|---|---|---|---|---|---|---|
| 271.919 | 1.88E-54 | 2 | 4 | 152092208 | 45047038 | rs41525550 | rs41497647 |
| 247.286 | 3.16E-49 | 16 | 18 | 68082045 | 66494570 | rs41511445 | rs41534148 |
| 50.4629 | 1.47E-08 | 8 | 23 | 99083471 | 69101723 | rs11776602 | rs5936809 |
| 49.6178 | 2.14E-08 | 2 | 5 | 148054025 | 109072457 | rs12473660 | rs6897334 |
| 48.07 | 4.22E-08 | 1 | 5 | 154126715 | 123388807 | rs1749405 | rs10463770 |
| 47.7293 | 4.90E-08 | 1 | 8 | 215419112 | 51015468 | rs17696625 | rs1552380 |
| 47.3635 | 5.74E-08 | 1 | 15 | 39927775 | 95905196 | rs538758 | rs12593275 |
| 47.015 | 6.69E-08 | 1 | 3 | 236849682 | 142760341 | rs9970821 | rs13084444 |
| 46.8823 | 7.09E-08 | 2 | 4 | 82858244 | 24686864 | rs7558379 | rs2024337 |
| 42.7404 | 4.26E-07 | 4 | 10 | 11477884 | 92201087 | rs2200819 | rs7914851 |
| 42.041 | 5.75E-07 | 3 | 16 | 170668093 | 2002066 | rs34585560 | rs8051877 |
| 40.7052 | 1.02E-06 | 7 | 11 | 2652585 | 40527074 | rs10950811 | rs7107769 |
| 36.5206 | 5.96E-06 | 4 | 14 | 12730820 | 66557382 | rs565867 | rs1950282 |
| 35.3225 | 9.81E-06 | 4 | 16 | 3480396 | 84745512 | rs3182 | rs1809844 |
| 33.0522 | 2.50E-05 | 4 | 23 | 141328479 | 22643958 | rs17005617 | rs3935725 |

Table 24: For HT dataset, reciprocally significant SNP pairs found by Potpourri RC1 - $k = 750$ and $\omega = 1.31623$ and multiple hypothesis testing threshold is $\sim 3.56E - 08$.

| **Chi-Sq.** | **P-value** | **SNP1 Chr** | **SNP2 Chr** | **SNP1 - Pos** | **SNP2 - Pos** | **SNP1 - rsID** | **SNP2 - rsID** |
|---|---|---|---|---|---|---|---|
| 271.919 | 1.88E-54 | 2 | 4 | 152092208 | 45047038 | rs41525550 | rs41497647 |
| 247.286 | 3.16E-49 | 16 | 18 | 68082045 | 66494570 | rs41511445 | rs41534148 |
| 126.035 | 8.93E-24 | 3 | 20 | 152638073 | 19899246 | rs41401648 | rs41384844 |
| 50.4629 | 1.47E-08 | 8 | 23 | 99083471 | 69101723 | rs11776602 | rs5936809 |
| 49.936 | 1.86E-08 | 2 | 15 | 148054025 | 23389643 | rs12473660 | rs4906728 |
| 48.4619 | 3.55E-08 | 1 | 15 | 228104209 | 97322652 | rs2780626 | rs2654981 |
| 48.07 | 4.22E-08 | 1 | 5 | 154126715 | 123388807 | rs1749405 | rs10463770 |
| 46.8996 | 7.03E-08 | 2 | 4 | 44335603 | 11470631 | rs12712916 | rs6448840 |
| 46.8823 | 7.09E-08 | 2 | 4 | 82858244 | 24686864 | rs7558379 | rs2024337 |
| 46.3411 | 8.97E-08 | 10 | 14 | 114154136 | 23294875 | rs11195949 | rs4982795 |
| 44.6297 | 1.89E-07 | 11 | 12 | 30401716 | 9004848 | rs489032 | rs11048264 |
| 44.5191 | 1.98E-07 | 8 | 14 | 62416109 | 46810794 | rs10100364 | rs8017252 |
| 42.7404 | 4.26E-07 | 4 | 10 | 11477884 | 92201087 | rs2200819 | rs7914851 |
| 42.678 | 4.37E-07 | 7 | 21 | 40327027 | 34860195 | rs12701816 | rs8130115 |
| 40.7052 | 1.02E-06 | 7 | 11 | 2652585 | 40527074 | rs10950811 | rs7107769 |
| 39.0851 | 2.03E-06 | 1 | 3 | 236061906 | 32305813 | rs4659819 | rs4484229 |
| 38.0789 | 3.10E-06 | 1 | 2 | 8144743 | 44328678 | rs17032281 | rs6712059 |
| 37.3478 | 4.21E-06 | 3 | 16 | 163650090 | 86119554 | rs6789378 | rs6540057 |
| 36.9287 | 5.02E-06 | 6 | 11 | 169803426 | 73686511 | rs9396992 | rs474280 |
| 36.5206 | 5.96E-06 | 4 | 14 | 12730820 | 66557382 | rs565867 | rs1950282 |
| 35.3225 | 9.81E-06 | 4 | 16 | 3480396 | 84745512 | rs3182 | rs1809844 |
| 34.4058 | 1.43E-05 | 3 | 20 | 73521155 | 30491788 | rs11128334 | rs6141721 |
| 34.2863 | 1.51E-05 | 12 | 18 | 110174962 | 76069383 | rs2078851 | rs9956207 |
| 33.0522 | 2.50E-05 | 4 | 23 | 141328479 | 22643958 | rs17005617 | rs3935725 |
| 32.8897 | 2.67E-05 | 3 | 3 | 9314644 | 39864600 | rs2648563 | rs930917 |

Table 25: For HT dataset, reciprocally significant SNP pairs found by Potpourri RC2 - $k = 750$ and multiple hypothesis testing threshold is $\sim 3.56E - 08$.

| **Chi-Sq.** | **P-value** | **SNP1 Chr** | **SNP2 Chr** | **SNP1 - Pos** | **SNP2 - Pos** | **SNP1 - rsID** | **SNP2 - rsID** |
|---|---|---|---|---|---|---|---|
| 271.919 | 1.88E-54 | 2 | 4 | 152092208 | 45047038 | rs41525550 | rs41497647 |
| 247.286 | 3.16E-49 | 16 | 18 | 68082045 | 66494570 | rs41511445 | rs41534148 |
| 53.1555 | 4.49E-09 | 21 | 23 | 45139519 | 39609316 | rs2838726 | rs3008916 |
| 50.4629 | 1.47E-08 | 8 | 23 | 99083471 | 69101723 | rs11776602 | rs5936809 |
| 49.7218 | 2.04E-08 | 4 | 10 | 11477195 | 49995061 | rs12645979 | rs11100990 |
| 49.6178 | 2.14E-08 | 2 | 5 | 148054025 | 109072457 | rs12473660 | rs6897334 |
| 48.4619 | 3.55E-08 | 1 | 15 | 228104209 | 97322652 | rs2780626 | rs2654981 |
| 48.07 | 4.22E-08 | 1 | 5 | 154126715 | 123388807 | rs1749405 | rs10463770 |
| 47.9794 | 4.39E-08 | 3 | 4 | 70722153 | 93811680 | rs4974301 | rs34905965 |
| 47.7293 | 4.90E-08 | 1 | 8 | 215419112 | 51015468 | rs17696625 | rs1552380 |
| 47.3635 | 5.74E-08 | 1 | 15 | 39927775 | 95905196 | rs538758 | rs12593275 |
| 47.0689 | 6.53E-08 | 2 | 10 | 79635544 | 50027795 | rs2685159 | rs2725190 |
| 47.015 | 6.69E-08 | 1 | 3 | 236849682 | 142760341 | rs9970821 | rs13084444 |
| 46.8823 | 7.09E-08 | 2 | 4 | 82858244 | 24686864 | rs7558379 | rs2024337 |
| 46.7924 | 7.37E-08 | 2 | 11 | 15922327 | 73693247 | rs2890496 | rs10898976 |
| 42.8173 | 4.12E-07 | 2 | 16 | 240105797 | 60338107 | rs7574895 | rs4494526 |
| 42.7404 | 4.26E-07 | 4 | 10 | 11477884 | 92201087 | rs2200819 | rs7914851 |
| 42.1849 | 5.41E-07 | 4 | 5 | 42084497 | 119187135 | rs6447159 | rs10793801 |
| 42.041 | 5.75E-07 | 3 | 16 | 170668093 | 2002066 | rs34585560 | rs8051877 |
| 40.7052 | 1.02E-06 | 7 | 11 | 2652585 | 40527074 | rs10950811 | rs7107769 |
| 38.6397 | 2.45E-06 | 5 | 12 | 65784598 | 56279757 | rs4700112 | rs812315 |
| 37.8283 | 3.44E-06 | 3 | 4 | 65476046 | 174112846 | rs9845819 | rs6825049 |
| 36.865 | 5.16E-06 | 7 | 22 | 147197951 | 44993808 | rs2373269 | rs4253754 |
| 36.5206 | 5.96E-06 | 4 | 14 | 12730820 | 66557382 | rs565867 | rs1950282 |
| 35.3225 | 9.81E-06 | 4 | 16 | 3480396 | 84745512 | rs3182 | rs1809844 |
| 34.9733 | 1.13E-05 | 7 | 10 | 2757211 | 21354589 | rs798500 | rs663661 |
| 34.6041 | 1.32E-05 | 1 | 20 | 145020771 | 18566017 | rs12129691 | rs6081267 |
| 33.0522 | 2.50E-05 | 4 | 23 | 141328479 | 22643958 | rs17005617 | rs3935725 |

Table 26: For HT dataset, reciprocally significant SNP pairs found by Potpourri RC3 - $k = 750$ and $\omega = 1.31623$ and multiple hypothesis testing threshold is $\sim 3.56E - 08$.

| Chi-Sq. | P-value | SNP1 Chr | SNP2 Chr | SNP1 - Pos | SNP2 - Pos | SNP1 - rsID | SNP2 - rsID |
|---|---|---|---|---|---|---|---|
| 271.919 | 1.88E-54 | 2 | 4 | 152092208 | 45047038 | rs41525550 | rs41497647 |
| 247.286 | 3.16E-49 | 16 | 18 | 68082045 | 66494570 | rs41511445 | rs41534148 |
| 126.035 | 8.93E-24 | 3 | 20 | 152638073 | 19899246 | rs41401648 | rs41384844 |
| 50.4629 | 1.47E-08 | 8 | 23 | 99083471 | 69101723 | rs11776602 | rs5936809 |
| 49.936 | 1.86E-08 | 2 | 15 | 148054025 | 23389643 | rs12473660 | rs4906728 |
| 48.7907 | 3.08E-08 | 3 | 12 | 2121505 | 98419467 | rs9310629 | rs924507 |
| 48.4619 | 3.55E-08 | 1 | 15 | 228104209 | 97322652 | rs2780626 | rs2654981 |
| 48.07 | 4.22E-08 | 1 | 5 | 154126715 | 123388807 | rs1749405 | rs10463770 |
| 46.8996 | 7.03E-08 | 2 | 4 | 44335603 | 11470631 | rs12712916 | rs6448840 |
| 46.8823 | 7.09E-08 | 2 | 4 | 82858244 | 24686864 | rs7558379 | rs2024337 |
| 46.3411 | 8.97E-08 | 10 | 14 | 114154136 | 23294875 | rs11195949 | rs4982795 |
| 46.0136 | 1.03E-07 | 9 | 11 | 116137576 | 129241367 | rs10739407 | rs10894158 |
| 45.393 | 1.35E-07 | 2 | 16 | 82388469 | 7657329 | rs11692865 | rs10492760 |
| 44.7803 | 1.77E-07 | 11 | 12 | 30401716 | 9038836 | rs489032 | rs1805721 |
| 44.5191 | 1.98E-07 | 8 | 14 | 62416109 | 46810794 | rs10100364 | rs8017252 |
| 42.8173 | 4.12E-07 | 2 | 16 | 240105797 | 60338107 | rs7574895 | rs4494526 |
| 42.7404 | 4.26E-07 | 4 | 10 | 11477884 | 92201087 | rs2200819 | rs7914851 |
| 42.678 | 4.37E-07 | 7 | 21 | 40327027 | 34860195 | rs12701816 | rs8130115 |
| 41.9307 | 6.03E-07 | 6 | 21 | 6195283 | 44473420 | rs3024378 | rs7278940 |
| 41.6951 | 6.67E-07 | 1 | 1 | 160329647 | 163962350 | rs1337062 | rs1913846 |
| 41.6111 | 6.91E-07 | 1 | 1 | 160329876 | 164000107 | rs1337061 | rs6426940 |
| 40.7052 | 1.02E-06 | 7 | 11 | 2652585 | 40527074 | rs10950811 | rs7107769 |
| 40.6265 | 1.05E-06 | 9 | 20 | 115176814 | 6299417 | rs1040989 | rs6076987 |
| 38.2952 | 2.83E-06 | 1 | 16 | 228535580 | 3548257 | rs7414930 | rs1859379 |
| 38.0789 | 3.10E-06 | 1 | 2 | 8144743 | 44328678 | rs17032281 | rs6712059 |
| 37.8283 | 3.44E-06 | 3 | 4 | 65476046 | 174112846 | rs9845819 | rs6825049 |
| 37.3478 | 4.21E-06 | 3 | 16 | 163650090 | 86119554 | rs6789378 | rs6540057 |
| 36.9287 | 5.02E-06 | 6 | 11 | 169803426 | 73686511 | rs9396992 | rs474280 |
| 36.865 | 5.16E-06 | 7 | 22 | 147197951 | 44993808 | rs2373269 | rs4253754 |
| 36.5206 | 5.96E-06 | 4 | 14 | 12730820 | 66557382 | rs565867 | rs1950282 |
| 35.3225 | 9.81E-06 | 4 | 16 | 3480396 | 84745512 | rs3182 | rs1809844 |
| 33.0522 | 2.50E-05 | 4 | 23 | 141328479 | 22643958 | rs17005617 | rs3935725 |
| 31.4484 | 4.80E-05 | 5 | 15 | 60625496 | 94618471 | rs4546329 | rs7182413 |

Table 27: For T2D dataset, reciprocally significant SNP pairs found by LINDEN. Multiple hypothesis testing threshold is $\sim 1.31E - 11$.

| Chi-Sq. | P-value | SNP1 Chr | SNP2 Chr | SNP1 - Pos | SNP2 - Pos | SNP1 - rsID | SNP2 - rsID |
|---|---|---|---|---|---|---|---|
| 219.613 | 2.26E-43 | 3 | 4 | 152638073 | 44958383 | rs41401648 | rs28576190 |
| 195.927 | 2.23E-38 | 1 | 6 | 156523434 | 3468258 | rs41362848 | rs41525244 |
| 145.539 | 8.00E-28 | 2 | 12 | 177023053 | 100660931 | rs41331247 | rs41402844 |
| 68.0659 | 5.45E-12 | 17 | 22 | 13840981 | 36004493 | rs9897978 | rs12170152 |
| 67.291 | 7.75E-12 | 4 | 8 | 63945084 | 72141558 | rs17672919 | rs17699685 |

Table 28: For BD dataset, reciprocally significant SNP pairs found by LINDEN. Multiple hypothesis testing threshold is $\sim 1.31E - 11$.

| **Chi-Sq.** | **P-value** | **SNP1 Chr** | **SNP2 Chr** | **SNP1 - Pos** | **SNP2 - Pos** | **SNP1 - rsID** | **SNP2 - rsID** |
|---|---|---|---|---|---|---|---|
| 797.766 | 3.10E-167 | 4 | 16 | 26983021 | 47086325 | rs41472746 | rs41334049 |
| 82.0237 | 8.88E-15 | 2 | 16 | 222817551 | 60341749 | rs12328631 | rs17822810 |
| 77.5211 | 7.12E-14 | 23 | 23 | 128164876 | 144648869 | rs4497124 | rs2748640 |
| 76.625 | 1.08E-13 | 23 | 23 | 86625117 | 144939805 | rs6624074 | rs2102122 |
| 75.864 | 1.53E-13 | 23 | 23 | 12484265 | 37956756 | rs6641046 | rs7883635 |
| 74.761 | 2.54E-13 | 23 | 23 | 121630018 | 141779863 | rs716209 | rs991072 |
| 73.304 | 4.96E-13 | 4 | 23 | 94723510 | 39230319 | rs10516922 | rs1318833 |
| 72.8267 | 6.17E-13 | 23 | 23 | 88167972 | 143538098 | rs5985083 | rs2815689 |
| 72.2251 | 8.13E-13 | 9 | 13 | 106459062 | 75157071 | rs7872725 | rs9530460 |
| 72.1629 | 8.37E-13 | 23 | 23 | 113067002 | 147338454 | rs5974183 | rs3860290 |
| 71.9181 | 9.36E-13 | 23 | 23 | 17469973 | 100850029 | rs1894579 | rs5951290 |
| 71.574 | 1.10E-12 | 23 | 23 | 50583279 | 77521778 | rs5915321 | rs321034 |
| 71.2936 | 1.25E-12 | 23 | 23 | 5167871 | 85395112 | rs7888448 | rs5968843 |
| 70.7603 | 1.59E-12 | 23 | 23 | 26751837 | 152929084 | rs5986678 | rs3027898 |
| 69.5221 | 2.80E-12 | 23 | 23 | 69141924 | 124523655 | rs6625561 | rs2097323 |
| 68.9297 | 3.67E-12 | 5 | 6 | 179993410 | 16526541 | rs10085109 | rs9370893 |
| 68.9089 | 3.71E-12 | 5 | 10 | 125902892 | 14799607 | rs730870 | rs10906750 |
| 68.7402 | 4.01E-12 | 23 | 23 | 77851450 | 141999744 | rs7052801 | rs6636902 |
| 68.6583 | 4.16E-12 | 23 | 23 | 82767162 | 111740955 | rs2506825 | rs17308025 |
| 68.2191 | 5.08E-12 | 10 | 17 | 122053346 | 42622984 | rs10886660 | rs11570441 |
| 67.9357 | 5.78E-12 | 23 | 23 | 21952813 | 95733494 | rs178685 | rs4969592 |
| 67.7647 | 6.25E-12 | 23 | 23 | 99654701 | 150568197 | rs720403 | rs5924659 |
| 67.5852 | 6.78E-12 | 3 | 23 | 171473104 | 97820677 | rs2140825 | rs2335828 |
| 67.3747 | 7.46E-12 | 12 | 23 | 126980427 | 43262688 | rs4761036 | rs4986513 |
| 67.2868 | 7.77E-12 | 23 | 23 | 118783580 | 147329420 | rs2782240 | rs6540370 |
| 67.2828 | 7.78E-12 | 23 | 23 | 112879733 | 144801782 | rs5929410 | rs12862591 |
| 67.1621 | 8.22E-12 | 23 | 23 | 28831428 | 87712529 | rs5985809 | rs6617715 |
| 67.1036 | 8.45E-12 | 2 | 14 | 19825202 | 75769285 | rs12464329 | rs935331 |
| 67.0841 | 8.52E-12 | 11 | 23 | 130238715 | 95901267 | rs6590517 | rs5949456 |
| 67.0655 | 8.59E-12 | 16 | 23 | 23584612 | 85837950 | rs35585 | rs5923597 |
| 66.7898 | 9.74E-12 | 5 | 23 | 137957669 | 26899255 | rs4835690 | rs6526586 |
| 66.5545 | 1.08E-11 | 23 | 23 | 117831542 | 141600993 | rs759144 | rs11096012 |
| 66.3656 | 1.18E-11 | 23 | 23 | 8611505 | 148523250 | rs5978943 | rs550402 |
| 66.3614 | 1.18E-11 | 11 | 12 | 106553206 | 2816231 | rs4754188 | rs4765999 |
| 66.3439 | 1.19E-11 | 2 | 23 | 98909640 | 117743282 | rs12467504 | rs2495630 |

Table 29: For HT dataset, reciprocally significant SNP pairs found by LINDEN. Multiple hypothesis testing threshold is $\sim 1.32E - 11$.

| **Chi-Sq.** | **P-value** | **SNP1 Chr** | **SNP2 Chr** | **SNP1 - Pos** | **SNP2 - Pos** | **SNP1 - rsID** | **SNP2 - rsID** |
|---|---|---|---|---|---|---|---|
| 593.828 | 2.46E-123 | 2 | 16 | 225162111 | 47086325 | rs41419450 | rs41334049 |
| 67.4735 | 7.14E-12 | 1 | 8 | 63117028 | 137411912 | rs1537496 | rs16905646 |

Table 30: Time performances of LINDEN, Potpourri-guided LINDEN and Potpourri on the T2D dataset.

| Method | | T2D CPU Time (log10) | T2D Run Time (hh:mm:ss) |
|---|---|---|---|
| **LINDEN** | | 4.54 | 01:04:53 |
| **Potpourri guided LINDEN** | $k$ = *500* | 3.73 | 00:17:39 |
| | $k$ = *750* | 3.67 | 00:16:50 |
| | $k$ = *1000* | 3.79 | 00:24:40 |
| | $k$ = *1500* | 3.85 | 00:28:18 |
| | $k$ = *2000* | 3.88 | 00:30:20 |
| **Potpourri** | $k$ = *500* | 3.69 | 00:14:54 |
| | $k$ = *750* | 3.77 | 00:19:16 |
| | $k$ = *1000* | 3.75 | 00:20:47 |
| | $k$ = *1500* | 3.84 | 00:22:45 |
| | $k$ = *2000* | 3.89 | 00:22:30 |

Table 31: Runtime performances of LINDEN, Potpourri-guided LINDEN and Potpourri on the BD dataset.

| Method | | BD CPU Time (log10) | BD Run Time (hh:mm:ss) |
|---|---|---|---|
| **LINDEN** | | 4.82 | 01:25:18 |
| **Potpourri guided LINDEN** | $k$ = *500* | 3.78 | 00:14:41 |
| | $k$ = *750* | 3.66 | 00:16:28 |
| | $k$ = *1000* | 3.85 | 00:26:25 |
| | $k$ = *1500* | 3.76 | 00:27:34 |
| | $k$ = *2000* | 3.82 | 00:31:41 |
| **Potpourri** | $k$ = *500* | 3.78 | 00:15:14 |
| | $k$ = *750* | 3.75 | 00:18:35 |
| | $k$ = *1000* | 3.78 | 00:19:24 |
| | $k$ = *1500* | 3.86 | 00:21:16 |
| | $k$ = *2000* | 3.91 | 00:22:46 |

Table 32: Runtime performances of LINDEN, Potpourri-guided LINDEN and Potpourri on the HT dataset.

| Method | | HT CPU Time (log10) | HT Run Time (hh:mm:ss) |
|---|---|---|---|
| **LINDEN** | | 4.76 | 01:00:09 |
| **Potpourri guided LINDEN** | $k$ = *500* | 3.79 | 00:14:43 |
| | $k$ = *750* | 3.66 | 00:17:26 |
| | $k$ = *1000* | 3.86 | 00:26:14 |
| | $k$ = *1500* | 3.77 | 00:29:24 |
| | $k$ = *2000* | 3.83 | 00:32:11 |
| **Potpourri** | $k$ = *500* | 3.69 | 00:18:46 |
| | $k$ = *750* | 3.68 | 00:19:13 |
| | $k$ = *1000* | 3.77 | 00:20:01 |
| | $k$ = *1500* | 3.85 | 00:22:31 |
| | $k$ = *2000* | 3.91 | 00:24:34 |
